# Harnessing Matrix Completion to Unify and Extend Viral Serology Studies

**DOI:** 10.1101/2021.08.29.458105

**Authors:** Tal Einav, Brian Cleary

**Author notes:** Authors contributed equally to this work.

## Abstract

The development of new vaccine strategies, as well as our understanding of key processes that shape viral evolution and host antibody repertoires, rely on measuring multiple antibody responses against large panels of viruses. Given the enormous diversity of circulating virus strains and antibody responses, exhaustive testing of all antibody-virus interactions is infeasible. Even within individual studies with limited panels, exhaustive testing is not always done, and there is no common framework for combining information across studies with partially overlapping panels, especially when the assay type or host species differ. Results from prior studies have demonstrated that virus-antibody interactions can be characterized in a vastly simpler and lower-dimensional space, suggesting that relatively few measurements could accurately predict unmeasured antibody-virus interactions. Here, we apply matrix completion to several of the largest-scale studies for both influenza and HIV-1. We explore how prediction accuracy evolves as the number of available measurements changes and approximate the number of additional measurements necessary in several highly incomplete datasets (suggesting ∼250,000 measurements could be saved). In addition, we show how the method can combine disparate datasets, even when the number of available measurements is below the theoretical limit that guarantees successful prediction. This approach can be readily generalized to other viruses or more broadly to other low-dimensional biological datasets.

**Significance:** One of the central problems in immunology is to characterize how our vast array of antibodies inhibits the diverse pathogens we encounter in our lives. In this work, we apply a well-studied mathematical technique called matrix completion that leverages patterns in partially-observed antibody-virus inhibition data to infer unmeasured interactions. We predict the results of tens of thousands of missing experiments in influenza and HIV-1 studies and quantify the expected error of our estimates. By harnessing matrix completion, future experiments could be designed that only collect a fraction of measurements, saving time and resources while maximizing the information gained.

## Introduction

The constant evolution of viruses such as influenza and the human immunodeficiency virus (HIV-1) leads to viral escape and degraded immunity. As a result, the influenza vaccine must be periodically reformulated to focus our antibody response against currently circulating strains (Petrova & Russell, 2017). In the context of HIV-1, antibody cocktails are necessary to counteract the vast diversity of viral strains within each host (Wagh et al., 2018). Thus, the ability to characterize the spectrum of viral variants against panels of antibodies or antisera is critical to the development of new vaccine strategies and successful viral control.

While thousands of new variants emerge each year, only a small fraction can be functionally characterized in terms of their effects upon the antibody repertoire. Assays quantifying binding, neutralization, or hemagglutination inhibition (HAI) are time- and resource-intensive, necessitating small antibody-virus panels that rarely exceed several dozen strains (Creanga et al., 2021; Georgiev et al., 2013; Kong et al., 2015; Li et al., 2012; Wrammert et al., 2011). Crucially, virus panels often differ between studies, making it difficult to translate the lessons learned in one context to predict new viral behavior.

Nonetheless, such small-scale studies have helped shed light on how viral infection or vaccination reshape the antibody response (Amanna et al., 2007; Arevalo et al., 2020; J. Lee et al., 2019; Thompson et al., 2016) as well as how the existing antibody repertoire constrains viral evolution (Dingens et al., 2019; J. M. Lee et al., 2019; Perelson et al., 2012). Moreover, an increasing number of large-scale studies have probed how our antibody response copes with the diverse array of viruses we encounter. For example, each year the influenza surveillance network surveys ∼100,000 viruses against ∼10 reference sera to determine whether an antigenically-distinct virus strain has emerged. Combining this surveillance data with insights from small-scale studies could increase the resolution of antibody-virus interactions and lead to improved vaccine selection (Petrova & Russell, 2017).

This ability to combine datasets is especially feasible in the context of antibody-virus interactions, where past studies have repeatedly shown that the data are low-dimensional (Lapedes & Farber, 2001; Ndifon, 2011). Strains from the same lineage often elicit similar antibody responses and can be grouped together into antigenic clusters (Smith et al., 2004). Different antibodies targeting the same epitopes, or sera from individuals with similar exposure histories, can show correlated activity across viral strains (Fonville et al., 2014; Kucharski et al., 2015). The profile of serum inhibition from an individual exposed to two antigenically distinct viruses can be expressed as a combination of profiles from two individuals, each exposed to one of the viruses (Carter et al., 2016). Mathematically, these features are consistent with a low-rank structure (Figure S1).

The theory of compressed sensing, and the related field of matrix completion, describe how such structure can be leveraged in the setting of under-sampled experiments (Candes & Recht, 2009; Cleary & Regev, 2020). These techniques have been extensively studied theoretically and applied in many settings from online recommender systems to magnetic resonance imaging (Candes & Tao, 2010). With matrix completion, the goal is to use a relatively small number of observations (individual matrix entries; here, corresponding to a virus-antibody or virus-serum pair) to identify low-rank features that can be used to infer missing values.

In the context of antigenic characterization, matrix completion has been used to refine the calculation of antigenic distance between strains (Cai et al., 2010; Ndifon, 2011). However, prior work has not sufficiently addressed questions of sample complexity (describing how the magnitude of errors will change based on the number of available measurements), which is critical for the successful design of under-sampled experiments. Moreover, while earlier studies have performed matrix completion on the data within a given study (what we will call *intra*-table matrix completion, Figure 1A), the limits of *inter*-table matrix completion are not clear (Figure 1B). It remains to be demonstrated to what extent partially overlapping studies conducted in varying geographic regions and points in time can be integrated; whether different types of studies (*e.g*., infection versus vaccination, or using fluorescence versus hemagglutination inhibition (Fonville et al., 2014; Nguyen Vinh et al., 2021) can be combined; and whether studies across species (particularly human versus ferret in the case of influenza) can successfully inform each other in the context of matrix completion. Successful inference in these tasks could vastly increase the amount of data used to study viral evolution or develop vaccination strategies, and could be applied to improve ongoing experimental design.

**Figure 1.**
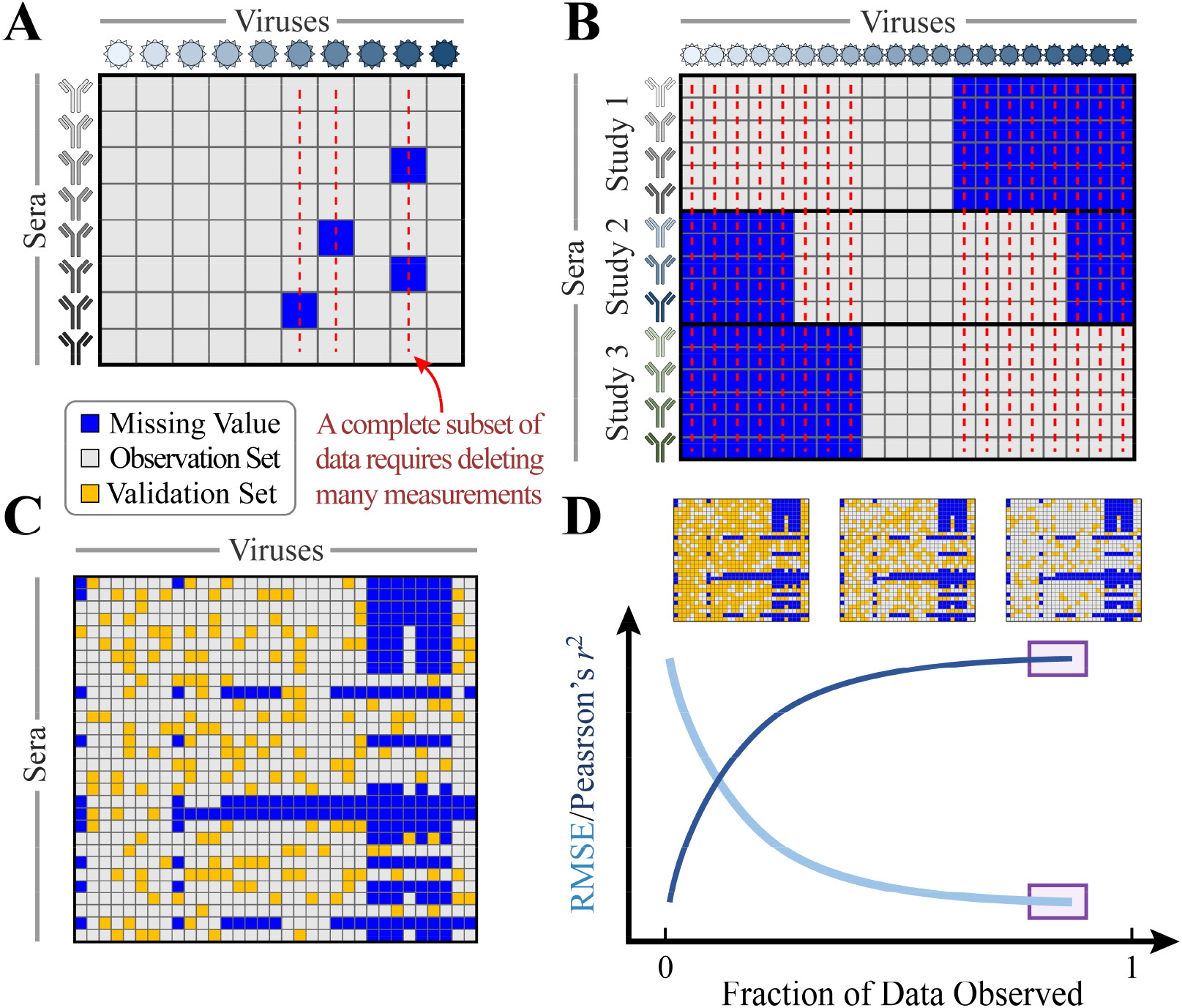
Harnessing low dimensionality to impute missing values. Schematic of hypothetical data showing missing antibody-virus measurements within a single study (A) or between multiple studies (B). Subsequent analyses without matrix completion may require entire columns/rows to be removed (red lines) to generate a complete subset of data. (C) Schematic of matrix completion on hypothetical data with missing values [blue squares]. To estimate the accuracy of the resulting completion for these missing values, we quantify how well an observation set [gray] can predict the values of a withheld validation set [gold]. (D) As the fraction of data observed increases, the root-mean-squared error (RMSE) decreases and Pearson’s correlation coefficient (*r*^2^) increases. The prediction accuracy for the missing values is estimated using the limit where the full data is used with no validation set [shown by the purple boxes].

Here, we develop a low-rank matrix completion framework that can analyze the increasing number of serological studies of the antibody response against viruses. We focus on three of the largest such datasets: (1) The Fonville influenza dataset consists of six studies (1 ferret study, 1 human infection study, and 4 human vaccination studies; collectively 1147 sera × 81 viruses), all measured using HAI (Fonville et al., 2014). (2) The Vinh influenza dataset contains one study (human infection study, 24,000 sera × 6 viruses) measured using HA (Nguyen Vinh et al., 2021). (3) The Catnap HIV-1 dataset contains two categories of data (monoclonal antibody data, 373 sera × 933 viruses; polyclonal serum data, 40 sera × 71 viruses) combining antibody-virus neutralization measurements from over one hundred studies (Yoon et al., 2015).

Such experiments commonly contain a subset of missing measurements. Ideally, subsequent analyses can ignore these missing values and use all available data — yet in some cases, such as quantifying the dimensionality of the antibody response using singular value decomposition, entire rows or columns must be removed to obtain a complete subset of data (Figure 1A, red lines). Moreover, measurements across multiple studies are difficult to combine since virus panels tend to only partially overlap, further constraining the number of measurements (Figure 1B). For example, only 9/81 viruses were included in all six Fonville studies, and 0/933 of the Catnap viruses were included across the full database.

In this work, rather than trimming down to a complete subset of data, we instead harness the low dimensionality of the immune response and extrapolate missing antibody-virus measurements both within and between studies. Mathematically, missing values are imputed by minimizing the nuclear norm of the antibody-virus data to create a complete and low-rank matrix. This framework can be implemented with fewer than 10 lines of code in Python or Mathematica (see Methods and the linked <u>GitHub repository</u>). In the following sections, we illustrate this approach using progressively more complex forms of matrix completion, creating a framework that naturally incorporates antibody-virus measurements from multiple studies to inform future efforts. In the final section, we show how datasets can be combined to predict hundreds of thousands of new measurements.

## Results

### Antibody-virus data are consistent with a low-rank structure

We first quantified how well a low rank approximation can characterize the influenza and HIV-1 datasets. Every dataset we evaluated is considerably low-rank in comparison to its naive dimensionality, with ∼95% of variance in entries explained by rank 6-23 approximations, depending on the dataset (Table S1). These low-rank approximations minimally shifted the existing measurements, with a root-mean-squared error (RMSE) ≤0.06 for the log_10_(titers) across all studies (*i.e*., un-logged titers are shifted by 10^0.06^=1.1-fold on average, representing near perfect fidelity).

These results hold for the nearly complete influenza studies (Fonville Studies 5-6; each 160 sera × 20 viruses and rank 8‒9) as well as the more incomplete studies (Fonville Studies 1-4; between 35‒324 sera × 57‒74 viruses and rank 6‒8) [Table S1]. Interestingly, the low rank of the Fonville data is similar to the number of antigenic clusters (*n*=14) in the combined panel of 81 viruses (Fonville et al., 2014), although substantial differences between viruses in the same antigenic cluster are observed in the completed matrix (Figure S2). The low rank result also holds for large tables with hundreds of antibodies and viruses, including the monoclonal antibody Catnap database (373 antibodies × 933 viruses and rank 23). The larger rank in the Catnap dataset reflects a greater number of viruses (∼12x more than the next largest study) in addition to the larger diversity between the viruses in HIV-1 versus influenza data.

This consistent, low-rank structure suggests that substantial information about missing entries may be contained in the already measured values. According to established theoretical results, if we wish to complete the missing entries of an *n*_1_×*n*_2_ matrix with minimal error, the number of available measurements (*m*) needs to be proportional to the rank (*r*) of the complete matrix and proportional to *n* log(*n*) (with *n*=max(*n*_1_,*n*_2_)) (Candes & Tao, 2010; Recht, 2011). The number of necessary measurements is also proportional to the coherence of the low-rank projection (*μ*), which quantifies how concentrated the low rank factors are on individual entries. Matrices with coherent factors — which in this context corresponds to individual antibodies or viruses with unique behavior that is orthogonal to the behavior of all other antibodies or viruses — require more measurements for successful recovery. Together, at least

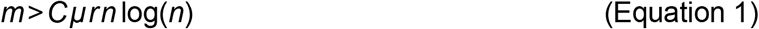

measurements are necessary, for some positive constant *C* that is context-specific (we approximate *C* below) (Candes & Tao, 2010).

While these results establish the sample complexity necessary for recovery with minimal error, there may be practical scenarios that deviate from some underlying assumptions of the theory. For example, the theoretical result assumes the available entries have been sampled uniformly at random from the matrix, but in reality, available data are likely to have been sampled in a biased (non-uniform) fashion and may require more measurements than the theoretical limit suggests. However, even when the number of samples is below this theoretical limit, the recovered values may nonetheless contain useful information about missing entries that might be used to effectively prioritize future experiments. Below we investigate how the error in completed entries evolves under increasing sample complexity, and under uniform and non-uniform sampling.

### Sample complexity under random uniform sampling

To quantify the accuracy of matrix completion within a single study using uniform random sampling (which we call intra-table completion), we split the available data in each study into observed measurements and a withheld validation set, and varied the fraction of entries withheld while completing the matrix using the observed entries as input (Figure 1C). As the observed fraction approaches 100% of available entries, we approximate the error of the truly missing values (shown schematically by the purple boxes in Figure 1D).

In the context of influenza, starting with 50% of measurements in Fonville Study 1 (the first of six studies within (Fonville et al., 2014)), we used matrix completion to impute the values of all missing and withheld titers (Figure 2A). We then quantified the accuracy of the withheld validation set, which exhibited a Pearson’s correlation coefficient of *r*^2^=0.8 (*i.e*., 80% variance explained) and an RMSE=0.4 for the log_10_(titers) (Figure 2B). To assess the theoretical limit for accurate recovery from Equation 1 (carried out below), we estimate the complexity of the completed data as the rank required to explain 95% of the variance, with rank=6 for the data shown (Figure 2C, Methods). This analysis is repeated for all six Fonville studies at varying fractions of observed data, with the accuracy of each completion quantified using Pearson’s correlation coefficient (*r*^2^) and the RMSE of the log_10_(titers) averaged over 10 runs to sample different subsets of data.

**Figure 2.**
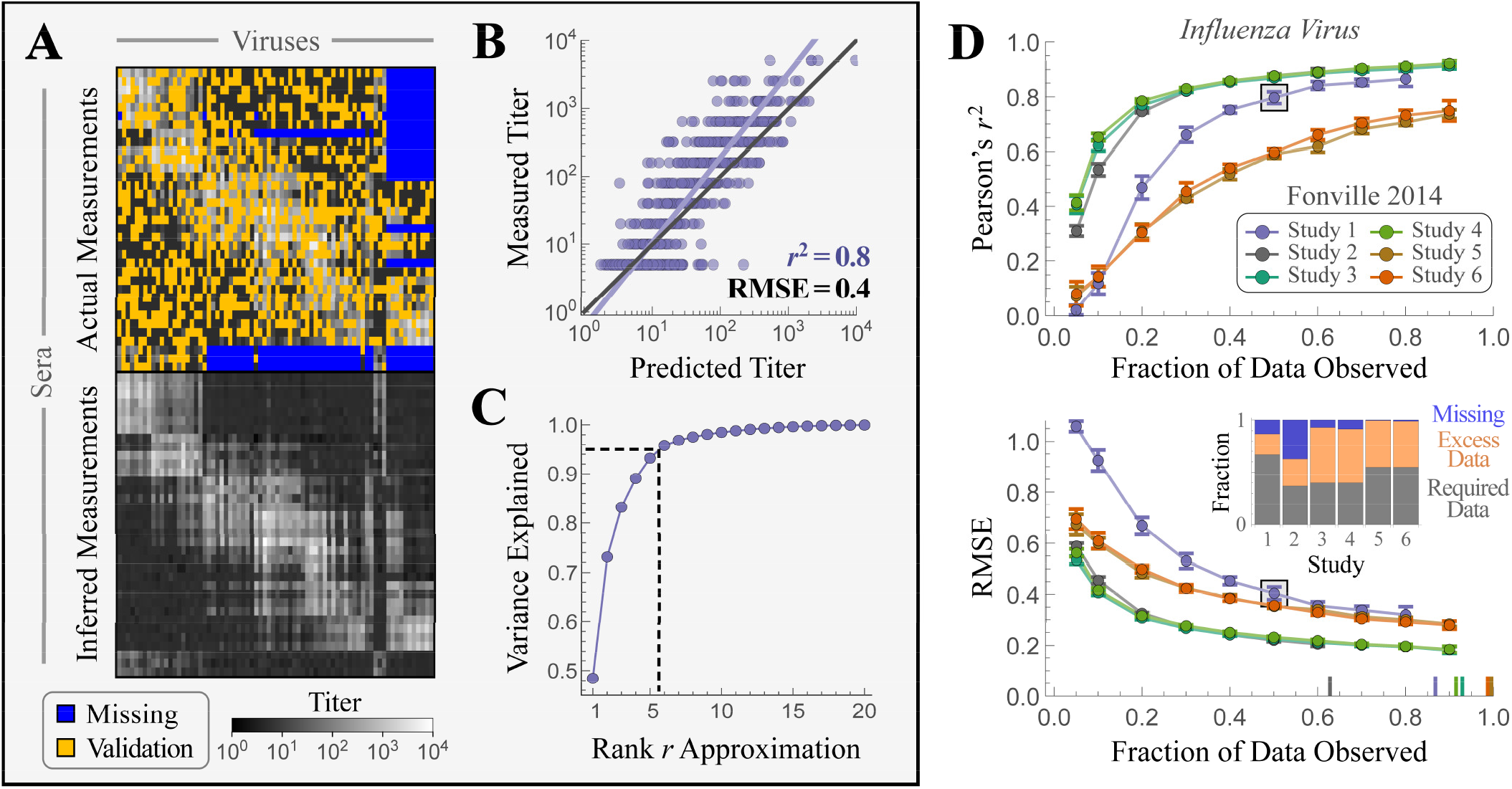
Completing antibody-virus measurements within influenza studies. (A-C) Low-rank matrix completion of Fonville Study 1 using 50% of all antibody-virus interactions. (A) Using available measurements [titers in grayscale] to infer the missing values [blue] and the withheld validation set [gold]. The top panel shows the input matrix and the bottom panel the completed matrix. (B) Predicted versus measured titers for the validation set withheld from the input matrix. (C) The variance of the completed matrix explained with a rank *r* approximation. The rank of a study is defined as the value *r* necessary to explain 95% of the variance (rank=6 for the data shown). (D) 10 iterations of intra-table completion at different fractions of observed data were performed for all six Fonville studies [individual lines], with the (mean±standard deviation) Pearson’s *r*^2^ and RMSE shown. Panels A-C represent 1 of 10 iterations at an observed fraction of 50% [boxed point on both plots]. The results for Studies 2-4 and Studies 5-6 are nearly overlapping. The markers on the *x*-axis of the RMSE plot denote the total fraction of antibody-virus interactions measured in each study (Table S1). Inset: The fraction of data in each study that is required for accurate matrix completion (gray), excess measurements that could have been inferred (orange), or missing measurements (blue).

For nearly complete studies with few missing values, we expect the final estimated error to approach the experimental accuracy of the assay, whereas the error for studies with a sizable fraction of missing values will depend on how the number of measurements compares to the bound discussed above (Equation 1). In the nearly complete tables (Fonville Studies 5 and 6), we indeed observe that the RMSE approaches 0.3 (in log_10_ units) (Figure 2D), comparable to the 10^0.3^=2-fold accuracy of the HAI assay (see Figure S8 of (Fonville et al., 2014)). As more entries are withheld, the RMSE increases, as expected (Figure 2D). The same trend is observed in the more incomplete influenza datasets, as well as the HIV-1 Catnap database (Figure 3), with the final estimated RMSE for the log_10_ titers across all tables varying from 0.2 in Fonville Studies 3-4 (90% of the full dataset available) to 0.8 in Catnap (13% available). Therefore, by varying the fraction of data observed, we can estimate both the error of the imputed titers and whether additional measurements will lead to markedly better predictions.

**Figure 3.**
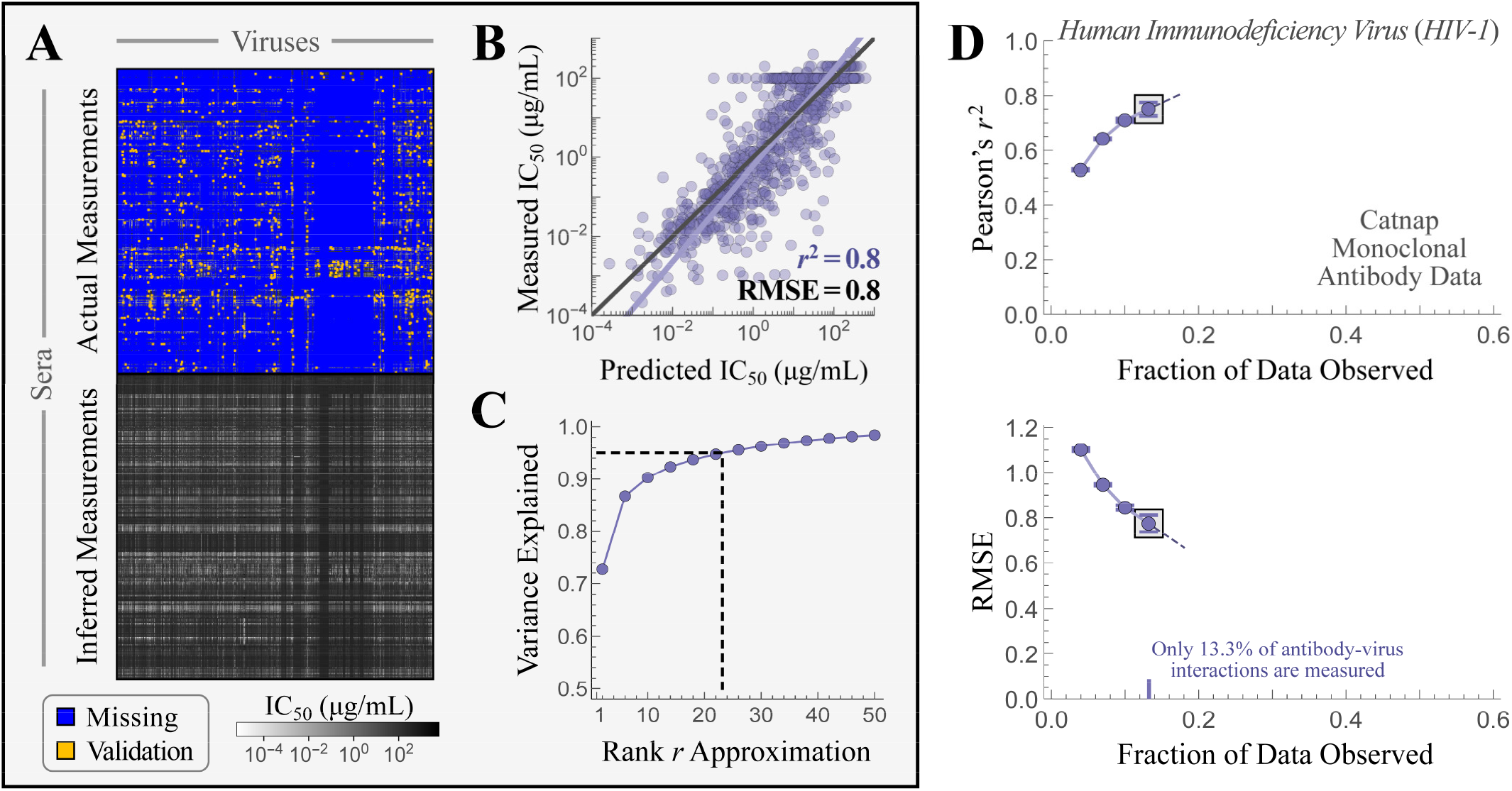
Analyzing a conglomeration of large-scale HIV-1 serology studies. (A-C) Low-rank matrix completion of HIV-1 monoclonal antibody data from Catnap using 13% of antibody-virus pairs as input and the remaining 0.3% of pairs for validation. (A) Using available measurements [titers in grayscale] to infer the missing values [blue] and the withheld validation set [gold]. The top panel shows the input matrix containing mostly missing values and the bottom panel shows the completed matrix. (B) Predicted versus measured titers for the antibody-virus measurements withheld in the input matrix. (C) The variance of the completed matrix explained with a rank *r* approximation. (D) 10 iterations of intra-table completion at different fractions of observed data were performed, with the (mean±standard deviation) Pearson’s *r*^2^ and RMSE shown. The last two points are linearly extrapolated to emphasize the expected change with additional measurements. The plots in Panel A represent 1 of 10 iterations at an observed fraction of 13% [boxed point on both plots].

Since all six Fonville studies impute the missing entries with an RMSE on the order of the experimental error, we can use these curves to determine how fewer measurements could produce equally successful completions. In brief, because the experimental error may vary by dataset and assay, we determine when the slope of each RMSE curve bottoms-out (when slope≥-0.5, see Methods). We find that in Fonville Study 1, 67% of measurements were needed (implying ∼1,000 measurements could have been “saved”); in Fonville Study 2, 37% were needed (∼12,000 saved measurements); in Studies 3 and 4, 40% were needed (∼20,000 saved measurements); and in Studies 5 and 6, 55% were needed (∼3,000 saved measurements) (inset of Figure 2D). The total across all studies, compared with measuring every value (including those missing in the published datasets), is ∼35,000 potential saved measurements out of a total of ∼60,000 possible measurements (58%).

We next examined intra-table completion in the context of HIV-1, where we used the more incomplete Catnap database to approximate the number of additional measurements necessary to achieve the lower bound RMSE. In monoclonal antibody studies, matrix completion using all available data led to an *r*^2^=0.8 and RMSE=0.8 for the missing values, with both metrics expected to rapidly improve with additional measurements (Figure 3). To approximate the minimum number of additional measurements necessary to achieve the lowest possible RMSE, we first use the available data in each Fonville study to estimate the coherence *μ* from Equation 1 (see Methods). We then use the number of necessary measurements empirically determined above, together with the theoretical lower bound in Equation 1, to find the value of the constant, *C*, in each Fonville study (Table S2), and average this value across the studies. Finally, given this constant, the size of the monoclonal antibody Catnap table, and estimates of its rank and coherence, we approximate the number of additional measurements necessary for successful completion (Table S2). This suggests that 27% of all possible measurements are necessary, implying that ∼48,000 of the ∼300,000 missing antibody-virus pairs need to be measured. We repeated this analysis for the polyclonal antibody Catnap data, which, like the Fonville tables, was sufficiently complete to get an empirical estimate of the fraction of measurements necessary for successful completion, and found good agreement between the empirical and theoretical results (44% and 41%, respectively; Table S2). We stress, however, that the theoretical estimates are a rough approximation, meant more to identify the scale for potential under-sampling, rather than attempting to define a specific point at which measurements should stop.

### Non-uniform sampling: Withholding a virus in inter-table completion

In the previous section, we examined the data in each study individually (*intra*-table completion), and the withheld validation set was uniformly sampled from all available measurements. In practice, missing or available measurements may not be spread uniformly across the matrix. For example, when combining data from different studies, which we call *inter*-table completion, missing measurements might appear in a more structured way (*e.g*., as blocks of missing values in Figure 1B). Relatedly, when predicting the results of experiments done in coming years using previously generated data, which we can evaluate in the Catnap database using the timestamps of each measurement, the available or missing data might be biased towards a few antibodies or viruses.

We first asked if inter-table completion can predict the inhibition of viruses that were not included in a given study. This form of completion is especially useful for rapidly evolving pathogens such as influenza or HIV-1, where studies published in the same year will often only have partially overlapping virus panels (Figure 1B).

We evaluated the accuracy of inter-table completion by merging all the Fonville studies, resulting in a matrix with 1147 serum samples and 81 viruses. For each of the 311 virus-study pairs with available data (Figure S3), we withheld from the merged matrix all measurements for that virus in that study, and asked whether the withheld values could be successfully inferred using matrix completion on the remaining data (Figure 4A, where the gold column show one virus withheld from Study 6). Since every virus had available measurements in at least two studies, the withheld virus was still functionally characterized, albeit in the contexts of a different study. We found that, on average, the imputed titers had an RMSE=0.3±0.2 (Figure 4B, gray bands), corresponding to roughly 2-fold error when predicting the behavior of a virus that is entirely absent from a given study.

**Figure 4.**
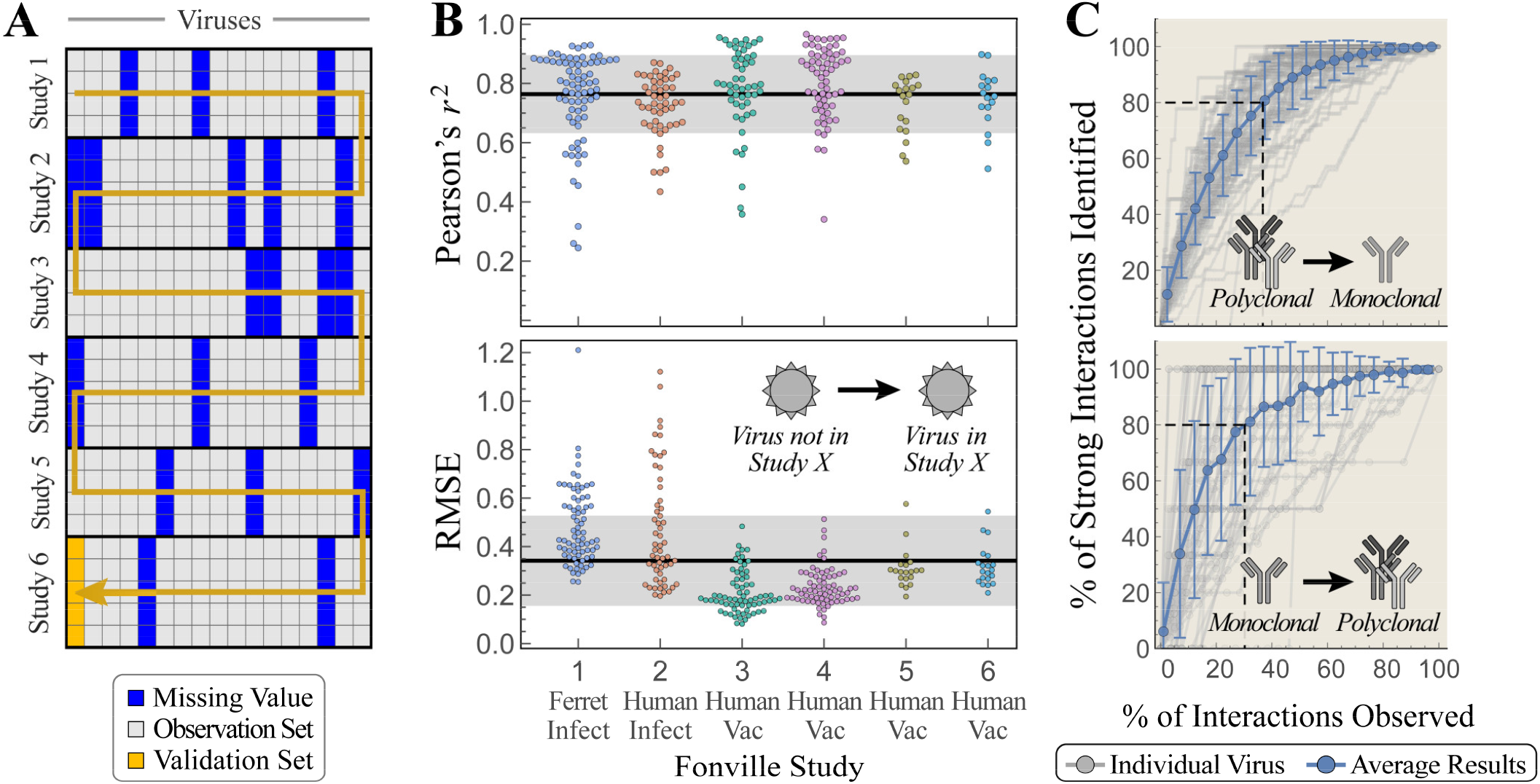
Predicting the behavior of viruses outside of a study. (A) Schematic of the overlapping virus panels across the six Fonville studies. For each virus *V*, we withhold *V*’s titers from one study [gold column], perform inter-table completion, and compare the predicted versus measured titers. The arrow represents all possible ways to remove a virus from a study, although viruses that have no measurements within a study [blue columns] are skipped. (B) One by one, each of the 81 Fonville viruses was withheld from one study (311 combinations, Figure S3). Swarm plots depict *r*^2^ [top] and RMSE [bottom] when withholding all possible viruses from each study. The mean±standard deviation is shown as gray bands. Viruses whose predicted or measured log_10_(titers) have standard deviation ≤0.2 were excluded from the *r*^2^ analysis since their correlation coefficient was artificially deflated (Methods). (C) Monoclonal antibody data and polyclonal serum data from Catnap against 71 shared viruses were combined. For each virus, all its measurements were withheld from either the monoclonal or polyclonal dataset and imputed via inter-table completion. These imputed values were ranked from most-to-least potent antibody interactions, and in that order the true measurements were revealed and characterized as strong interactions provided either the IC_50_<0.1 μg/mL (for antibody measurements) or ID_50_>1,000 (for sera). The gray lines represent results for individual viruses, and show how quickly the strong interactions are identified when using these matrix completion results, with the average over all viruses shown in blue. The fraction of measurements required to identify 80% of the potent antibodies is emphasized by the dashed lines.

While the majority of inter-table completions led to accurate predictions, some virus-study pairs showed a large RMSE (Figure 4B). Inter-table completion performed best for Fonville studies #3-4 and #5-6. Both pairs of studies used the same virus panel, and each contained human vaccination data collected in sequential years. Therefore, these studies represent the ideal scenario to apply inter-table completion, since virus trends in one study should be highly informative of the trends in the other study. Indeed, removing any virus from these four studies led to low reconstruction error (RMSE=0.2±0.1 for Studies 3-4 and RMSE=0.3±0.1 for Studies 5-6). To quantify how much effort could have been saved in each pair of studies, we completely removed *multiple* viruses and computed the average RMSE for the imputed values (Figure S4). In each case, removing half the viruses from one study only increased the average RMSE by 30%. This suggests that when two contemporary studies with similar experimental designs have partially overlapping virus panels, the two studies can be merged and accurately matrix completed.

On the other hand, Fonville Study 1 (ferret study) and Study 2 (human infection study) did not have a paired study with an identical virus panel and experimental design (Figure S5). Hence, these two studies represent a harder but more common scenario for inter-table completion. While many of the viruses in these first two studies had accurate inter-table completions (average RMSE=0.5±0.2 for both studies), there were some extreme outliers. For example, the hardest virus to predict (A/Auckland/20/2003) appeared in Studies 1, 3, and 4; it exhibited a large RMSE=1.2 when removed from Study 1 but a small RMSE=0.3 when excluded from Study 3 or 4. Moreover, restricting the inter-table completions to a subset of studies — such as imputing human infection data (Study 2) using human vaccination data (Studies 3-6) without ferret data (Study 1) — leads to marginally worse completions, and more generally, including additional and varied datasets leads to better completions (Figure S5). Indeed, when using all data, 99/131=75% of the inter-table completions for viruses in Study 1 or 2 had an RMSE<0.6, demonstrating that behavior of multiple viruses can be inferred across disparate studies. These results suggest that inter-table completion should improve as more datasets are combined, providing a data-driven framework to unify diverse datasets and extend their predictions. Moreover, the following sections demonstrate that inter-table completion can expedite experiments even in the low-data limit where the imputed values are less accurate.

### Non-uniform sampling: Combining monoclonal and polyclonal antibody data

We next evaluated a different form of inter-table completion using monoclonal antibody data to impute polyclonal serum measurements (and vice versa) in the Catnap database. While both studies measure the amount of antibodies required to neutralize a virus by 50%, monoclonal antibodies represent a single, defined antibody whereas polyclonal serum contain an unknown mixture of antibodies. Hence monoclonal antibodies are measured in terms of the 50% inhibitory concentration (IC_50_ in μg/mL units; lower values represent a more potent antibody) while sera are measured by the dilution necessary to achieve 50% neutralization (ID_50_ in dilution units; higher values represent a more potent serum). This more difficult case of inter-table completion must combine these disparate types of data as well as compensate for greater incompleteness of the data, with a far larger number of monoclonal antibody measurements (373 antibodies × 933 viruses) than polyclonal data (40 sera × 71 viruses; each of these viruses is also present in the monoclonal data).

Consistent with these expectations, we observed higher inter-table RMSEs than in Fonville studies where all the data were of the same type. The average error when withholding viruses from the monoclonal study was RMSE=1.2±0.3 while the average error when withholding from the polyclonal study was RMSE=0.7±0.3, implying an absolute prediction error of roughly 16-fold or 5-fold, respectively. However, given that the correlation between the imputed and missing entries was also somewhat high (average *r*^2^ of 0.47 and 0.49 when withholding from the monoclonal and polyclonal studies, respectively), the imputed data may contain useful information. For example, these predictions could differentiate between the weakest and strongest of the missing interactions.

Hence, despite the larger magnitude of errors, we hypothesized that inter-table completion could identify the most potent antibodies or sera. For each virus-study pair in the merged monoclonal-polyclonal Catnap data, we sorted the inferred entries for the withheld virus from most neutralizing to least neutralizing. We then asked whether prioritizing future experiments in this order would identify highly inhibitory antibodies (IC_50_<0.1 μg/mL for monoclonal; ID_50_>1,000 dilutions for polyclonal data) more effectively than performing future experiments in a random order. Averaging across viruses withheld from the monoclonal study, only the first 36% of prioritized experiments were necessary to identify 80% of highly inhibitory antibodies (Figure 4C). Similarly, the top 30% of prioritized experiments were necessary to identify 80% of the top hits in the polyclonal study (Figure 4C). Both cases improve upon a random search, which would require 80% of measurements to be carried out to find 80% of the most inhibitory interactions. Thus, even when matrix completion predictions have large errors, they nonetheless contain actionable information for the efficient design of future experiments.

### Non-uniform sampling: Using timestamps to predict Catnap measurements across time

We next examined how well existing measurements can predict future experiments. To that end, we turned to the dozens of studies comprising the monoclonal antibody-virus interactions in the Catnap database that together create the largest matrix we analyzed (350,000 interactions; Figure 3A). Each measurement is timestamped with the year it was deposited, enabling us to quantify how well past measurements could predict future data.

As noted from our intra-table completion analysis above, 87% of entries are missing from this monoclonal antibody data, and the imputed titers exhibited a relatively high estimated RMSE=0.8 (signifying 6-fold error). This large error not only arises from the large fraction of missing entries but also from the considerable divergence between the HIV-1 viruses in the Catnap database, which differ by 200 amino acids on average, far exceeding the average 30 amino acid difference between the Fonville influenza viruses. It is easier to matrix complete a virus’s titers when a similar virus exists in the dataset, and conversely, the accuracy of matrix completion decreases as the distance to the nearest virus increases (Figure S6A). In the context of HIV-1 where all viruses are highly diverse, the baseline matrix completion error tends to be higher, but it also depends less on sequence similarity (Figure S6B). Yet although the matrix completion results had a large RMSE for the missing values, they also exhibited a large *r*^2^=0.7. As with the monoclonal-polyclonal data examined above, this large *r*^2^ implies that the completed titers can accurately distinguish which of the missing values represent very weak or very potent interactions. Said another way, 6-fold error is more than sufficient to differentiate strong and weak antibody interactions, given that IC_50_ values span 6 orders of magnitude from 10^−4^‒10^2^ μg/mL.

To assess this approach, we examined how well *N* experiments, prioritized by matrix completion, could identify weak antibody-virus interactions (indicating viral escape) or the strongest interactions (representing a potent immune response), with *N* varying from 1 to the total number of experiments performed. We used the timestamp associated with each Catnap measurement and considered data taken before a given year (*e.g*., in 2017 or before) to predict the missing values that were subsequently measured (*e.g*., in 2018 or later; Figure 5A). In doing so, we only consider the antibodies and viruses measured both before 2017 and after 2018 (in the example of Figure 5A, we ignore the antibody in the right-most column and the virus in the bottom row). We then ordered the predictions from strongest-to-weakest (Figure 5B) and determined the number of strong interactions (defined as an IC_50_≤0.1 μg/mL) identified by *N* experiments.

**Figure 5.**
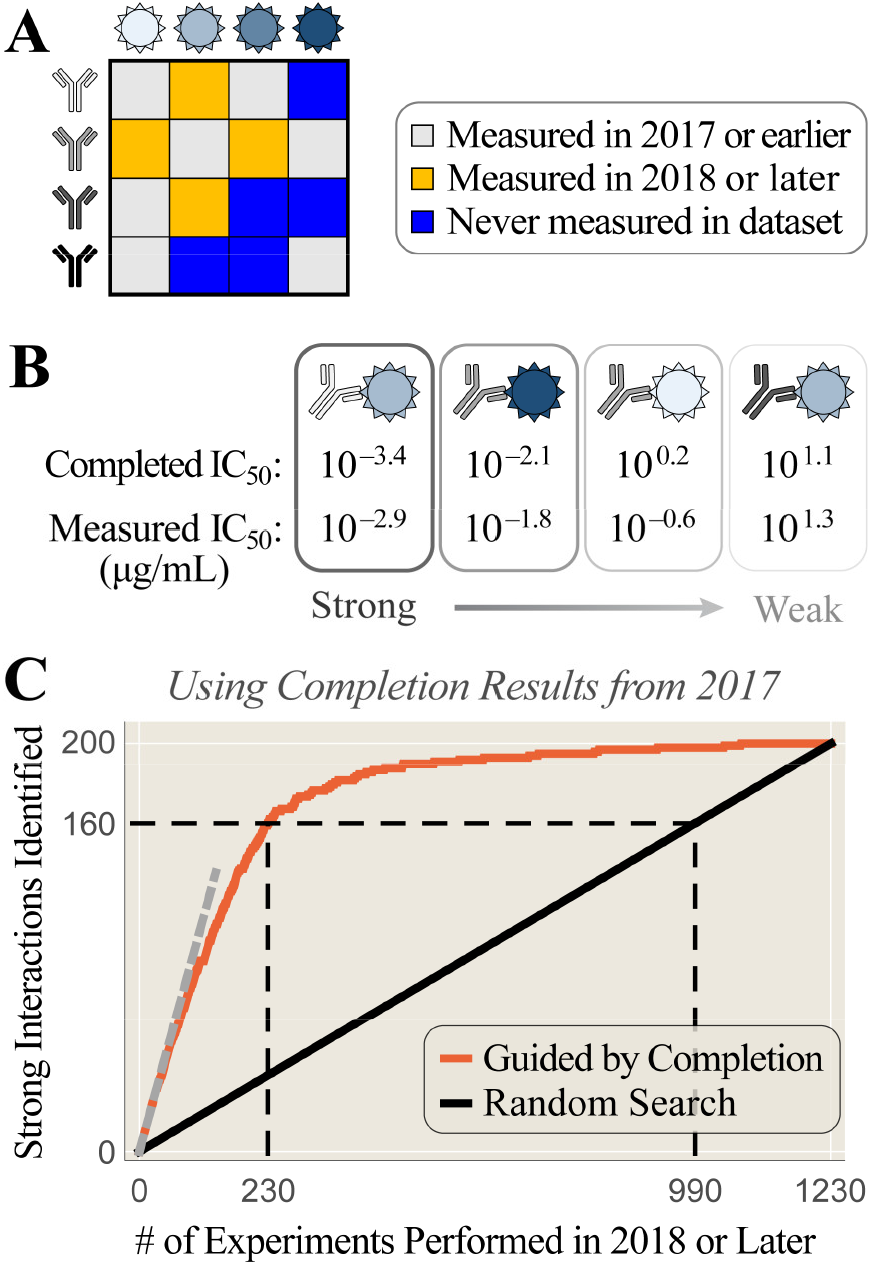
Prioritizing high-potency measurements in future studies. (A) Matrix completion using all Catnap monoclonal antibody-virus entries from 2017 or before [gray] but withholding measurements from 2018 or later [gold]. (B) The imputed IC_50_s are ordered from the strongest-to-weakest (first row of values). Note that smaller IC_50_s represent more potent antibody-virus interactions. We compare these predictions with the measurements made in 2018 or later (second row of values). (C) By carrying out *N* experiments (*x*-axis) ordered from the strongest-to-weakest imputed IC_50_s, the number of strong antibody-virus interactions discovered after 2018 (red curve) increases faster than a random search (black line). The black dashed line shows how many experiments are necessary to identify 80% of all strong interactions under both search schemes. The gray dotted line (*y*=*x*) represents perfect predictions for the strong interactions.

When using matrix completion to predict the most potent antibody-virus interactions, only 260/1230=20% of measurements were required to identify 160/200=80% of the strong interactions found in 2018 or later (Figure 5C). In contrast, searching through the antibody-virus pairs at random would have required 980 measurements on average before 80% of the strong interactions could have been found, necessitating 4x more work. A similar trend was found when searching for the weakest antibody-virus interactions or when analyzing the Catnap data before/after a different year (Figure S7). In this manner, matrix completion can sift through the wealth of available data to find new antibody-virus interactions with especially high or low titers.

### Applying inter-table completion to extend a small virus panel measured against 24,000 sera

Lastly, we applied inter-table completion to a recent serological study to generate hundreds of thousands of predictions that can guide future experiments. Specifically, we combine measurements from the Fonville 2014 influenza study described above (1147 sera × 81 viruses) with another large-scale influenza study by Vinh *et al*. (24,000 sera × 6 viruses; all six viruses have equivalent strains in the Fonville panel, as described in the Methods) (Fonville et al., 2014; Nguyen Vinh et al., 2021). For each virus in the Fonville panel whose behavior can be extrapolated to the Vinh sera, we generate 24,000 data points.

To ascertain which viruses can be accurately extrapolated, we first analyzed the 6 viruses in the Vinh study and repeated our earlier procedure, withholding measurements in the Vinh study for one virus at a time (Figure 6A). For this process, we used the completed Fonville data to fill in missing measurements against these six viruses, thereby leveraging all Fonville sera to inform the Vinh predictions. The predicted titers of the four viruses isolated between 2005-2011 exhibited a strong correlation (*r*^2^≥0.8) with the withheld values, and could effectively identify the potent antibody responses (Figure 6B, Figure S8). For all four strains, ≥80% of potent interactions would have been discovered by testing only 20% of sera sorted by matrix completion. In contrast, the two viruses isolated in 2003 and 1968 exhibited a weaker correlation and worse discovery rate (Figure 6B, Figure S8). Therefore, we restricted our application of inter-table completions to the 17 Fonville viruses isolated between 2005-2011. We combined the Vinh and Fonville measurements of the 6 viruses in the Vinh panel, together with Fonville measurements of one of the 17 additional viruses, and predicted the measurements for this additional virus against the 24,000 Vinh sera (Figure 6A, blue squares). We visualize the completed matrix by highlighting the 20% most potent sera for each of the Vinh and Fonville viruses (colored or black bars in Figure 6C; white bars represent the remaining 80% of sera).

**Figure 6.**
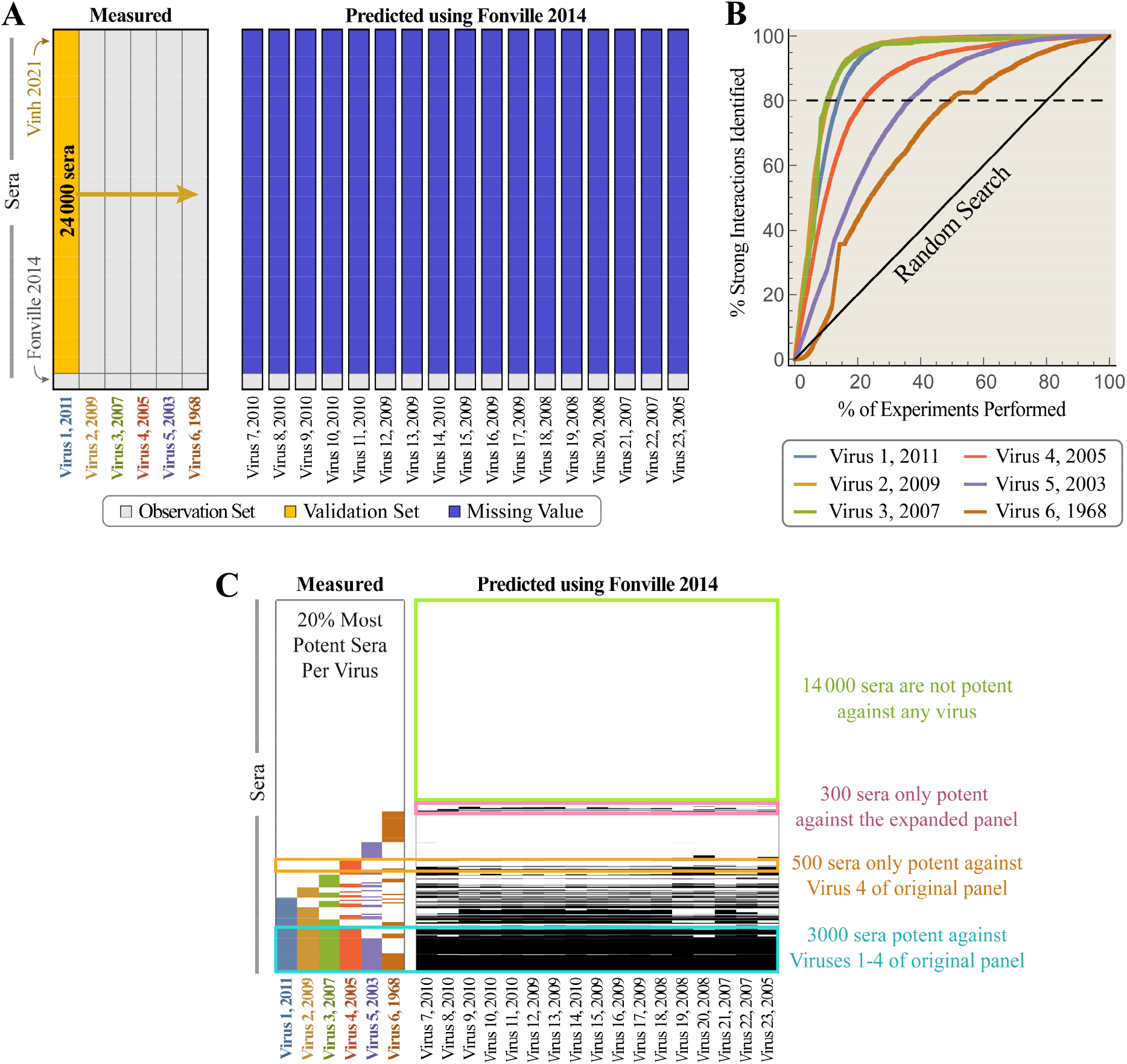
Predicting an expanded virus panel using inter-table completion. (A) Combining data from the two influenza studies Vinh 2021 (24,000 sera × 6 viruses) and Fonville 2014 (1147 sera × 81 viruses) (Fonville et al., 2014; Nguyen Vinh et al., 2021). 6 of the Fonville viruses are equivalent to the Vinh strains, while 17 other Fonville viruses were isolated between 2005-2011 and used to extrapolate the Vinh responses (Methods). [*Left*] Each of the six viruses is individually withheld and predicted via matrix completion. [*Right*] The complete Fonville and Vinh data predict how these 24,000 sera would interact with one of 17 additional Fonville viruses. (B) When one of the Vinh viruses is withheld [gold column], its imputed ID_50_s are ordered from the strongest-to-weakest. After carrying out a fraction of these prioritized experiments (*x*-axis), the number of strong antibody-virus interactions (ID_50_>1,000) discovered is shown for each virus. (C) For each virus, the 20% of sera with the strongest predicted interactions are shown as either colored or black bars (white bars represent the 80% of weaker interactions). The sera are first ordered by their interaction against Virus 1 [potent sera shown at the bottom-left in blue], then Virus 2 [the bottom gold segment shows sera potent against Virus 1 and 2; the short gold segment above that denotes sera only potent against Virus 2], then Virus 3, and so on.

As a sanity check, we find that 3000 sera that were uniformly potent against the 4 Vinh viruses isolated between 2005-2011 were also predicted to be highly potent against the Fonville viruses from this same time period [Figure 6C, cyan box]. Very few of these sera are predicted to inhibit 0-1 of the Fonville viruses (10 sera), and the majority of sera are predicted to inhibit ≥10 Fonville viruses (2700 sera).

The expanded virus panel further enables us to detect inhibition patterns that cannot be discerned from the Vinh data alone. For example, a set of 300 sera were outside the top 20% of potent interactions for all six Vinh viruses, yet they were in the top 20% for at least one Fonville strain [Figure 6C, pink box]. These sera exhibited a wide range of distinct inhibition profiles, ranging from potently inhibiting just 1 of the Fonville viruses (70 sera) to ≥10 viruses (80 sera). In this way, sera with specific inhibition profiles of interest against the extended virus panel — from the exceptionally broad to especially narrow — can be picked out from the 24,000 sera for further study.

## Discussion

The low dimensionality of antibody-virus data provides the mathematical foundation to fill in missing entries in a partially observed table of binding, neutralization, or HAI measurements. This same principle underlies the field of compressed sensing, where a sparse signal can reconstruct a complete image, enabling dramatically faster data acquisition. In the context of immunology, low dimensionality enables us to readily combine existing studies to infer missing measurements with high accuracy or, in cases of severe data limitation, rationally prioritize a minimal set of experiments to identify important virus-antibody interactions. Given the increasing number of large-scale serological studies, the power of data-driven techniques such as matrix completion will continue to grow, offering a simple framework that can unify diverse datasets and vastly increase the amount of available data.

We suggest several ways to harness matrix completion and facilitate future serological studies. First, quantifying how a large panel of antibody responses inhibits a large number of viruses does not require measuring each interaction. If the experiment can be performed in several rounds (*e.g*., randomly selecting 20% of the unmeasured antibody-virus pairs each time), then the error of imputing the missing values can be calculated after each round (Figures 2D). When this error becomes comparable to the error of the assay (typically 2-fold for HAI or neutralization measurements), or when the error bottoms-out, the remaining values can be filled in via matrix completion. In the context of the Fonville 2014 studies, this amounts to saving tens of thousands of measurements (corresponding to 50% fewer experiments).

Second, when only a small number of viruses or sera will be screened (Harvey et al., 2016; Nguyen Vinh et al., 2021), the few viruses or sera chosen should be as functionally orthogonal to one another as possible to maximize the information gained. The ability to successfully matrix complete with only a fraction of measurements would suggest that the viruses are functionally similar, and a more diverse virus panel could be chosen.

Beyond these applications of matrix completion on a single dataset (*intra*-table completion), utilizing completion across datasets (*inter*-table completion) remains largely unexplored. One of the simplest implementations is to merge datasets and infer measurements for viruses that were entirely missing from a study (Figure 1B). This method of inter-table completion consistently worked between pairs of human vaccination studies carried out in consecutive years (Fonville Studies 3-4 and 5-6, Figure 4B), presenting a simple framework to combine contemporary studies and greatly magnify the amount of data. Predictably, this approach was not always successful when inferring human (or ferret) infection data from human vaccination data or vice versa, as differences in ferret and human immune systems likely make the structure of these datasets more dissimilar than other Fonville studies we compared (Fonville Studies 1 and 2 in Figure 4B, Figure S5). Unlike with intra-table completions, where a fraction of the data can be withheld to quantify the expected error of the imputed values, there is currently no analogous method to quantify the expected error from inter-table completion. Hence, a fraction of interactions needs to be measured to verify the results of such matrix completions.

Even in cases where the error of imputed values is unacceptably large (for example, if the majority of measurements are missing or if the diversity of viruses/antibodies results in a large coherence), inter-table completion can nevertheless pinpoint the strongest or weakest interactions that are often of greatest interest. For example, while only 16% of all 1230 HIV-1 antibody-virus measurements from 2018-2020 had a strong IC_50_≤0.1 μg/mL, by matrix completing all data from 2017 and earlier, we determined a minimal set of 260 experiments that identified 80% of these strong IC_50_ values (Figure 5C). In addition, we demonstrated that monoclonal and polyclonal HIV-1 data against a panel of identical viruses could be combined to identify the strongest neutralization measurements, even though the composition of antibodies within these sera is not known (Figure 4C). Thus, if a new viral variant emerges and is rapidly tested against a panel of sera, matrix completion can determine which previously-characterized monoclonal antibodies potently inhibit the virus. Finally, we showed how combining datasets enables us to peer into the antibody responses at higher resolution by characterizing an antibody response against an expanded set of viruses (Figure 6). In short, inter-table completion can rapidly sift through all missing antibody-virus combinations and prioritize efforts for groups seeking the strongest or weakest interactions.

Unlike earlier applications of matrix completion using multidimensional scaling or alternating gradient descent (Cai et al., 2010; Lapedes & Farber, 2001; Ndifon, 2011; Smith et al., 2004), our approach of minimizing the nuclear norm does not require a dataset’s rank to be specified. Hence, it is straightforward to adapt this method to different studies and different viruses without needing to refit hyperparameters. For example, we used the same code to analyze the six influenza studies (with rank between 6‒9) and the HIV-1 Catnap dataset containing hundreds of studies (with rank 23). Further refinements to our approach could incorporate side information such as virus sequence (Radhakrishnan et al., 2021) or use tensor factorization to decompose higher-order data (Liu & Moitra, 2020).

Matrix completion can harness the low-rank behavior of antibody-virus interactions to greatly increase the amount of available data. In light of these results, it is worth considering how much data we actually have. Even when the viruses or antibodies characterized across multiple studies only partially overlap, patterns seen in one context can help inform other experiments. Indeed, from the vantage of matrix completion, differences in antibody and virus panels between experiments becomes an asset, enabling us to explore a greater fraction of the immune landscape and extrapolate the behavior of novel antibody-virus pairs.

## Methods

### Availability of Code and Results

The matrix-completed results for the Fonville and Catnap datasets are included in the supplementary files. We also include the code to perform matrix completion in both Mathematica and Python (https://github.com/cleary-lab/Inhibition-Neutralization-Completion). In Python, nuclear norm minimization takes fewer than 10 lines of code with the cvxpy package (Agrawal et al., 2018; Diamond & Boyd, 2016):

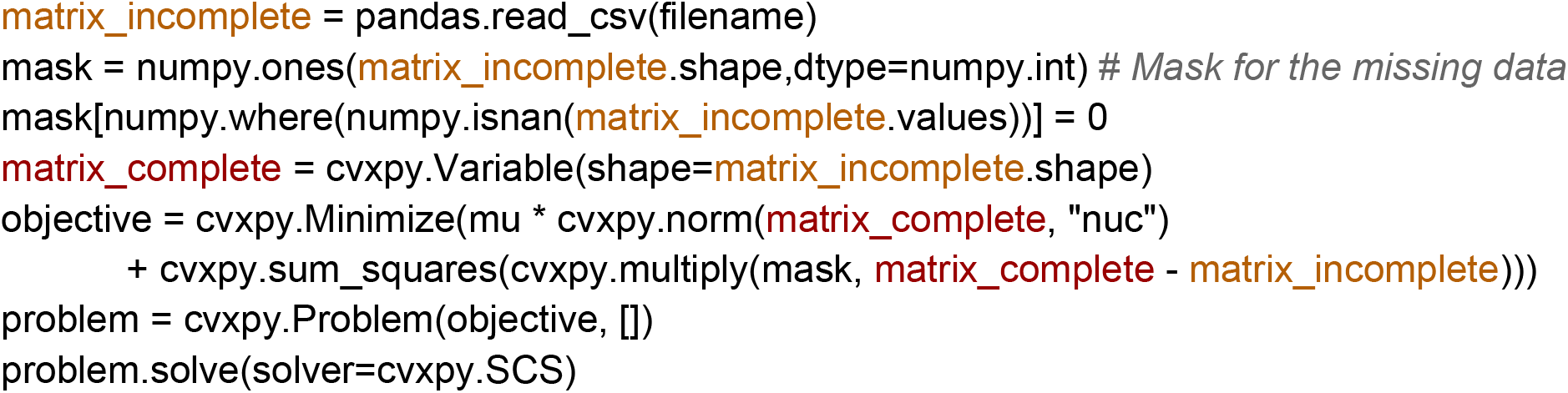

### Influenza: The Fonville 2014 Dataset

We gathered antibody-virus HAI measurements from the supplementary information of Fonville 2014 that contains the following six studies:

- Study 1 [Fonville Table S1]: Ferret study; 35 sera measured against 74 viruses
- Study 2 [Fonville Table S3]: Human infection study from 2007-2012; 324 sera measured against 57 viruses
- Study 3 [Fonville Table S5]: Human vaccination study from 1997; 212 sera measured against 70 viruses
- Study 4 [Fonville Table S6]: Human vaccination study from 1998; 256 sera measured against 70 viruses
- Study 5 [Fonville Table S13]: Human vaccination study from 2009; 160 sera measured against 20 viruses
- Study 6 [Fonville Table S14]: Human vaccination study from 2010; 160 sera measured against 20 viruses

Each serum sample only appeared in a single study, whereas all viruses were included in at least two studies. Cumulatively, the six studies contained 1147 distinct serum samples and 81 distinct viruses, with the distribution of viruses across the six studies shown in Figure S3. The raw data annotated with the virus names is included in the GitHub repository associated with this paper.

### Influenza: The Vinh 2021 Dataset

The Vinh dataset measures 24,000 sera against the following six H3N2 viruses:

- Virus 1 = A/Victoria/361/2011
- Virus 2 = A/Victoria/210/2009
- Virus 3 = A/Brisbane/10/2007
- Virus 4 = A/Wisconsin/67/2005
- Virus 5 = A/Wyoming/3/2003
- Virus 6 = A/Aichi/2/1968

The Fonville virus panel contains Vinh viruses #1, 3, 4, and 5, as well as two nearly equivalent strains to virus #2 (full name=A/Hanoi/EL201/2009, short name=HN201/2009, 7 amino acids away from A/Victoria/210/2009) and virus #6 (full name=A/Bilthoven/16190/1968, short name=BI/16190/68, 3 amino acids different from A/Victoria/210/2009). For our analysis, we associate the data from these two nearly equivalent strains. The Vinh panel also includes five H1N1 viruses, but we ignore those strains because the Fonville virus panel only contains H3N2 viruses.

### Intra-Table Matrix Completion

Consider a table of measurements *M* where entry *M*_*jk*_ represents the interaction between antibody (or serum) *j* and virus *k*, with larger values always corresponding to a more potent antibody that strongly inhibits a virus. *M*_*jk*_ either represents the hemagglutination inhibition (HAI) titer from the Fonville influenza study or the inverse of the 50% inhibitory concentration (1/IC_50_) for monoclonal HIV-1 antibodies from Catnap. Thus, potent antibody/serum interactions are always denoted by larger values.

Influenza HAI is measured using progressive 2-fold dilutions, so that titers can equal “<10”, 10, 20, …, 1280, or “≥1280.” We first replaced lower or upper bounds (“<10”→5 and “≥1280”→2560) and then took log_10_ of all titers. To prevent the error from large values from artificially dominating the completion algorithm, we subtracted the minimum value *m*=min_*jk*_(*M*_*jk*_) from all entries in *M*, performed matrix compilation, and then added *m* back to obtain the complete matrix with log titers (to transform to the original un-logged titers, the matrix completed value can be exponentiated by 10). We note that as an alternative, the mean of *M* could be subtracted to center the data in log-space; in this work, we chose to subtract the minimum titer because it results in slightly better predictions of the larger titer values (which are often the most important measurements).

The goal of matrix completion is to find a filled-in low-rank matrix 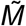 such that 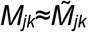 for all pairs (*j,k*) that were measured. Note these two criteria are at odds with one another, since maintaining exact fidelity with the data may result in a matrix 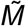 with larger rank. We control the relative importance of these two goals with a single hyperparameter λ described below.

Although minimizing matrix rank is NP-hard, minimizing the convex envelope of the rank (the nuclear norm ||*M*||⋆) is computationally efficient. Hence, the matrix *M*∈ ℝ ^*n*×*m*^ is completed as

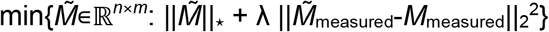

where 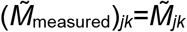 whenever entry (*j,k*) was measured and 0 otherwise, with an analogous definition for *M*_measured_. The first term, 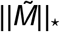, minimizes the nuclear norm (and hence the rank) of 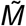, whereas the second term 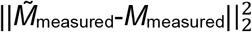 is a quadratic penalty for deviating from the measured values (the prefactor λ is discussed below). Such minimizations have a known tendency to shrink values towards zero, so that the best-fit line of the measured vs predicted titers has a slightly larger slope than the perfect experiment-theory match (purple vs black lines in the scatterplots of Figure 2A and 3A). For the influenza serum titers in Figure 2 where the log_10_(titers) are matrix completed, the predicted values tend to be smaller than the measured titers; for the HIV-1 Catnap antibody data where the -log_10_(IC_50_ values) are matrix completed, the predicted values tend to be greater than the measured titers because of the negation (see the *Catnap Database* Methods section).

The hyperparameter λ dictates the relative importance of the two terms. Large values of λ would indicate strong trust in the measured values and would result in higher-rank completions that remain highly faithful to the data. Smaller values of λ would be appropriate when the data is noisy and will result in a lower-rank matrix that can deviate from the measurements. In this work, we set λ=1 in all analyses. With this value, over 95% of the completed Fonville titers were within 10% of the measured values, keeping high fidelity with the experimental values.

Because the predicted and measured titers span multiple decades and are presented on log-log plots, both Pearson’s *r*^2^ and RMSE were computed on the log_10_ data. The RMSE can be recast into un-logged titers by exponentiating by 10 (*e.g*., an RMSE of 0.2 in log_10_ space becomes a 10^0.2^=1.6-fold error in the titer). The dimensionality of the data was computed on the titers mean-centered in log space, *M*_centered_ = log_10_(*M*) - mean(log_10_(*M*)) to prevent a change in the units (*e.g*., measuring an IC_50_ in μg/mL versus Molar) from affecting the dimensionality. The dimensionality *r* of *M*_centered_ is defined as the minimum number of singular values of *M*_centered_ that explain 95% of its variance (the minimum *r* such that 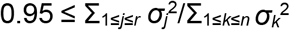).

### Inter-Table Matrix Completion

Computationally, inter-table completion works identically to intra-table completion, except that virus and serum panels must first be combined. Note that some of the Fonville viruses had different capitalization or punctuation in different tables (*e.g*., “HN196_2009” and “JO/33/94” in Fonville Table S1 and their counterparts “HN196/2009” and “A/JOHANNESBURG/33/94” in Fonville Table S3) that are unified in our supplementary data files. Any virus that was not present in a study was defined as missing for all sera in that study (*e.g*., “A/AUCKLAND/20/2003” was contained in Studies 1, 3, and 4, so when combining all Fonville data its titers were defined as “Missing” against sera in Studies 2, 5, and 6).

In the few cases when all predicted or measured log_10_(titers) withheld from a dataset have nearly identical values, the correlation coefficient *r*^2^ is artificially deflated. For example, if five (*predicted, measured*) titers for a virus were log_10_ of {(10,10), (10,10), (10,10), (10,20), (20,10)}, then any linear dependence is masked by the small range of the predictions and measurements. This leads to a small Pearson’s *r*^2^=0.06, even though the predictions were perfect for the first three points and only 2-fold off for the final two points. In contrast, when measurements span a wide dynamic range with this same error distribution, such as log_10_ of {(10,10), (20,20), (40,40), (80,160), (160,80)}, their resulting Pearson’s *r*^2^=0.81, matching the typical intuition for this metric. Hence, when all predicted or measured log_10_(titers) withheld had a standard deviation ≤0.2, we excluded such cases from the Pearson’s *r*^2^ analysis in Figures 4B, S3, and S5. We note that such cases only occurred for *inter*-table completion for viruses that showed minimal HAI against all sera (Figure 4B) and never occurred for *intra*-table completion (Figures 2 or 3).

When inter-table completing between the Fonville and Vinh datasets (Figure 6), we first matrix completed the Fonville data, then trimmed it to either the Vinh virus panel (6 viruses) or the extended virus panel (6+17 viruses), and then inter-table completed the Vinh measurements. This first matrix completion step is important, because many Fonville sera were not measured against all six Vinh viruses, and hence these sera would have been trimmed away without contributing to the Vinh completions. Performing this first matrix completion harnesses the full array of measurements in the Fonville dataset to make maximally informed predictions about the Vinh viruses.

When combining measurements from different studies (*e.g*., from the Fonville and Vinh studies), the accuracy of matrix completion improved when we *separately* mean-centered both datasets in log-space (*i.e*., divide all titers by the mean of the measured values). This can adjust for different protocols across labs or for systematic shifts in the data. Moreover, the accuracy improved when only a single additional virus was predicted (*i.e*., predicting the single withheld Vinh virus without the 17 additional Fonville strains). Thus, when we predicted the additional Fonville strains, we predicted the titers of one at a time.

### Time and Memory Usage

The nuclear norm described in this work scales polynomially with the size of the matrix. While the time required will vary with the dimensions of the matrix, a handy rule of thumb is that a matrix with 100 viruses and 100 sera takes ∼1 second to complete. Below are some benchmarks for the time and memory usage of matrix completion running Python on a high-performance computing machine:

- (2000 sera)×(6 viruses): 0.5 hours, 5 GB memory
- (2000 sera)×(80 viruses): 0.5 hours, 6 GB memory
- (6000 sera)×(6 viruses): 8 hours, 30 GB memory
- (6000 sera)×(80 viruses): 11 hours, 32 GB memory

The supplementary Mathematica notebook uses a robust PCA method (Candes et al., 2011) that yields nearly identical results to nuclear norm minimization, but which is over 100x faster. Below are these same benchmarks for the time and memory usage running Mathematica on a standard laptop:

- (2000 sera)×(6 viruses): 3 seconds, 7 GB memory
- (2000 sera)×(80 viruses): 30 seconds, 70 GB memory
- (6000 sera)×(6 viruses): 10 seconds, 20 GB memory
- (6000 sera)×(80 viruses): 50 seconds, 240 GB memory

### HIV-1: The Catnap Dataset

The complete list of Catnap antibody-virus measurements is updated each month on the Downloads page (https://www.hiv.lanl.gov/components/sequence/HIV/neutralization/download_db.comp; click the file labeled as “*Assay*”). We split each antibody-virus pair into one of four categories based on the antibody name and the IC_50_ and ID_50_ values:

1. *Polyclonal sera* (*n*=40) did not have IC_50_ values but instead had ID_50_ values (the dilution at which a virus was 50% neutralized).
2. *Antibody mixtures* (*n*=44) had a “+” in their antibody names to distinguish the monoclonal antibodies in each mixture.
3. *Multispecific antibodies* (*n*=50) had an “/” in their antibody name to denote the different monoclonal antibodies from which they were composed.
4. *Monoclonal antibodies* (*n*=373) contained all remaining entries.

For simplicity, our Catnap analysis was restricted to the monoclonal antibodies (category #4 above) and polyclonal sera (category #1). We replaced values bounded from above by half the value of the bound and values bounded from below by double their value (*e.g*., “<20”→10, “>40”→80), and when multiple antibody-virus measurements were available we took their geometric mean. We note that these assumptions, especially our treatment of the bounded values, may influence the matrix completion results given that the IC_50_ values in Catnap range from 10^−5^ μg/mL to 344 μg/mL, while the upper bounds varied between 10^−3^ and 10^0^.

The only minor difference between the Catnap analysis and the Fonville analysis accounted for the fact that potent antibodies have *low* IC_50_ values in Catnap whereas potent sera have a *high* HAI titer in Fonville. Therefore, we completed the matrix of -log_10_(IC_50_ values) from Catnap monoclonal antibody data, whereas we completed the matrix log_10_(HAI titers) for Fonville and log_10_(ID_50_) for Catnap polyclonal serum data. Note that in all cases, we represented the weak antibody/sera by smaller values in the matrix, which is important for the stability of the completion algorithm (because nuclear norm minimization pulls values towards 0).

### Number of Measurements Required for Accurate Matrix Completion

When missing values are randomly sampled across a matrix for intra-table completion, the theoretical bound in Equation 1 quantifies the number of measurements (*m*) required to accurately reconstruct a matrix based on the number of antibodies or viruses *n* (equal to max(# of antibodies, # of viruses)), the matrix rank *r* (here, determined as the rank required to explain 95% of the variance of the completed matrix), the coherence *μ* (calculated from the complete matrix as explained below), and a constant *C* (inferred from the Fonville RMSE curves where *m* can be determined empirically). Together, these values quantify how difficult it is to complete a given dataset.

To compute *μ* for all data within a study (Candes & Recht, 2009; Ndifon, 2011), we take all rows/columns of the data matrix with ≥50% of entries measured and calculate the completed matrix *M* whose dimensions we denote by *n*_1_×*n*_2_. (Note that using the rows/columns with few measurements leads to unstable results for highly incomplete datasets such as Catnap and are hence dropped.) We compute the rank *r* of *M* as the minimum number of singular values of *M* that explains 95% of its variance. Using the singular value decomposition *M*=*UΣV*^T^, we form the orthogonal projection onto *M* using the first *r* eigenvectors of *U* or *V*. For example, *P*_*U*_=Σ_1≤*j*≤*r*_ *u*_*j*_*u*_*j*_^T^ where *u*_*j*_ is the *j*^th^ column of *U*, with an analogous expression for *V*. The coherence of *M* is given by

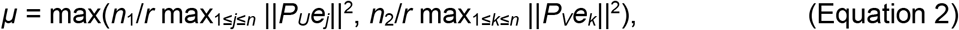

where *e*_*j*_ and *e*_*k*_ represent standard basis elements, and with values shown in Table S2.

To compute the value of the constant *C* in Fonville studies, we find the value of *m* at which the slope of RMSE curves in Figure 2 is greater than –0.5 (indicating the sufficient number of measurements for that study). We then solve for *C* in Equation 1, to obtain a separate value for each study (Table S2). Below, we use the averaged value, *C*=0.18.

To approximate when the Catnap RMSE and *r*^2^ curves in Figures 2B and 3B will flatten out, we utilize the theoretical bound in Equation (1) that a matrix can be accurately completed when the number of (randomly observed) measurements is greater than *Cμrn* log(*n*), where *n*=933 is the larger dimension of the Catnap matrix and *r*=23 equals the rank of the completed matrix (Candes & Recht, 2009). To validate this result, we apply this same approach to the Catnap polyclonal sera dataset and predict that the RMSE curve will bottom-out when 41% of measurements are observed. Since the polyclonal serum dataset has approximately half of all serum-virus interactions measured, we can directly test this prediction by performing intra-table completion using different fractions of observed data — in doing so, we find that the RMSE curves bottoms-out when 44% of measurements are observed, close to our predicted value.

## Acknowledgements

We would like to thank Jesse Bloom, John Huddleston, Heidi Klumpe, Richard Neher, and Armita Nourmohammad for their input on this manuscript. Tal Einav is a Damon Runyon Fellow supported by the Damon Runyon Cancer Research Foundation (DRQ 01-20). Brian Cleary was supported by a Merkin Institute Fellowship.

## Author Contributions

T.E. and B.C. conducted the research and wrote the paper.

## Competing Interests

The authors declare no competing interests.

## SI Figures and Tables

**Table S1.**
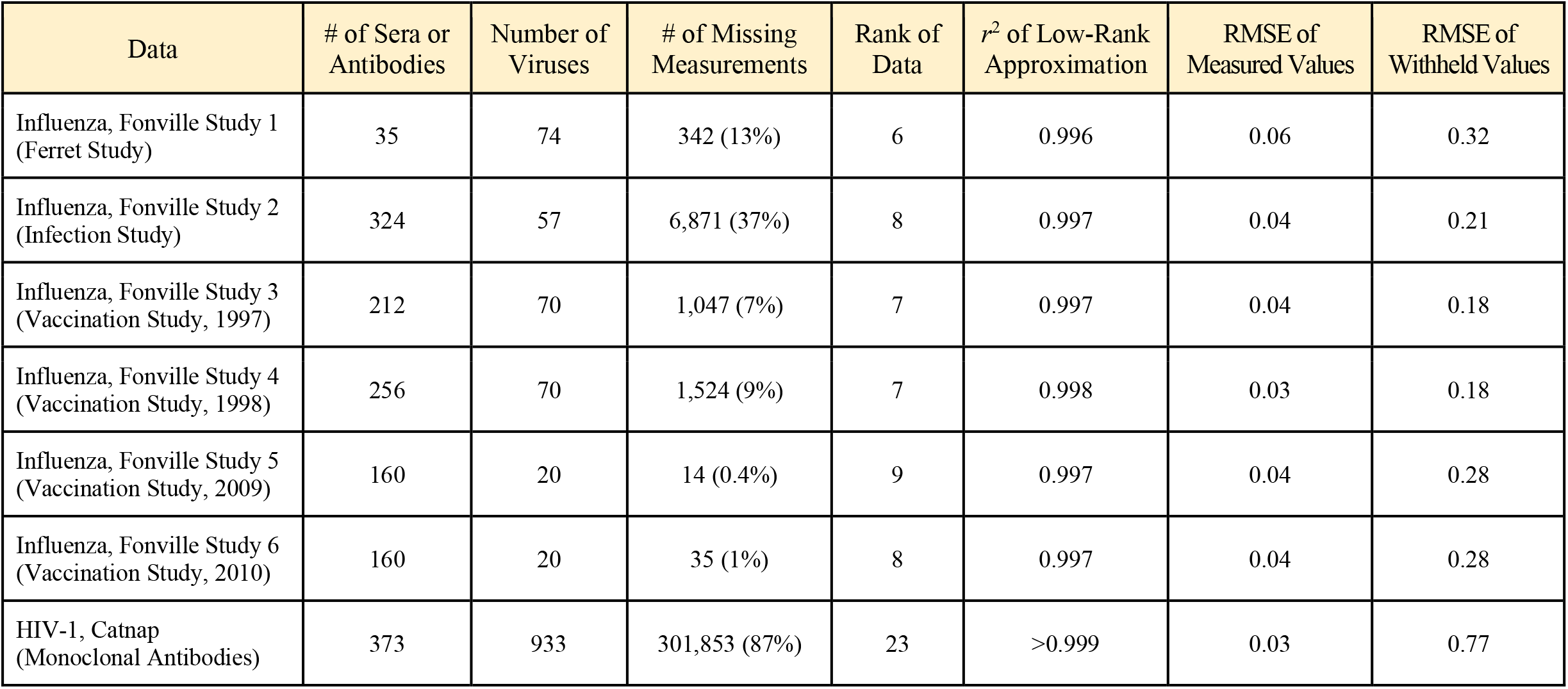
Datasets analyzed in this work. The size and number of missing entries in each dataset is shown. The rank of the data is defined as the number of singular values required to explain 95% of the variance of the low-rank completed matrix. The *r*^2^ of low-rank approximation shows the correlation between log_10_ of the measured values versus the low-rank complete matrix, with large values of ≈1 signifying that matrix completion alters the measured values very minimally. In the limit where the nearly the entire dataset is observed with only a few values withheld, the RMSE of the measured values quantifies the difference between the log_10_(measured titers) and their log_10_(low-rank approximation), while the RMSE of the withheld values estimates the accuracy of the log_10_(missing values) as the error of the withheld validation set.

| Data | # of Sera or Antibodies | Number of Viruses | # of Missing Measurements | Rank of Data | $r^2$ of Low-Rank Approximation | RMSE of Measured Values | RMSE of Withheld Values |
| --- | --- | --- | --- | --- | --- | --- | --- |
| Influenza, Fonville Study 1 (Ferret Study) | 35 | 74 | 342 (13%) | 6 | 0.996 | 0.06 | 0.32 |
| Influenza, Fonville Study 2 (Infection Study) | 324 | 57 | 6,871 (37%) | 8 | 0.997 | 0.04 | 0.21 |
| Influenza, Fonville Study 3 (Vaccination Study, 1997) | 212 | 70 | 1,047 (7%) | 7 | 0.997 | 0.04 | 0.18 |
| Influenza, Fonville Study 4 (Vaccination Study, 1998) | 256 | 70 | 1,524 (9%) | 7 | 0.998 | 0.03 | 0.18 |
| Influenza, Fonville Study 5 (Vaccination Study, 2009) | 160 | 20 | 14 (0.4%) | 9 | 0.997 | 0.04 | 0.28 |
| Influenza, Fonville Study 6 (Vaccination Study, 2010) | 160 | 20 | 35 (1%) | 8 | 0.997 | 0.04 | 0.28 |
| HIV-1, Catnap (Monoclonal Antibodies) | 373 | 933 | 301,853 (87%) | 23 | >0.999 | 0.03 | 0.77 |

**Table S2.**
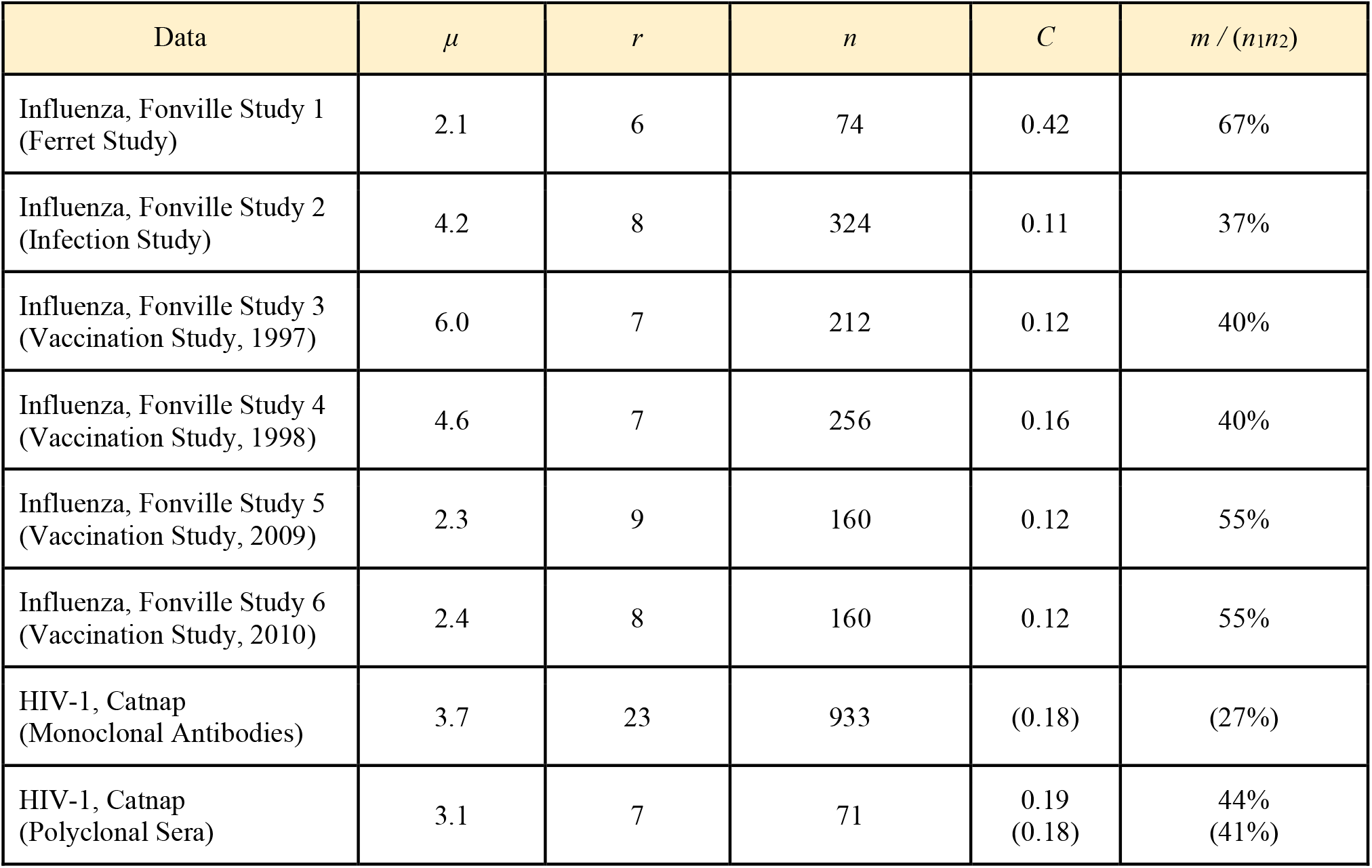
Number of measurements needed to accurately reconstruct missing data. The theoretical lower bound, *m* > *C μ r n* log(*n*), implies that accurate imputation via matrix completion requires at least *m* measurements in an *n*_1_×*n*_2_ matrix. *μ* represents the coherence of the matrix (Candes & Recht, 2009), *r* its rank, *n* the larger of the matrix length or width, and *C* is a constant that is determined from the RMSE curves of the Fonville studies. The average value of *C* is then used to determine at what fraction of observed data the Catnap RMSE curves will bottom-out (right-most column). The average value of the constant and the implied fraction of measurements necessary in Catnap studies are shown in parentheses. The Catnap polyclonal serum data is sufficiently complete to determine when the RMSE bottoms-out, providing a direct test for this approach.

| Data | $\mu$ | $r$ | $n$ | $C$ | $m / (n_1 n_2)$ |
| --- | --- | --- | --- | --- | --- |
| Influenza, Fonville Study 1 (Ferret Study) | 2.1 | 6 | 74 | 0.42 | 67% |
| Influenza, Fonville Study 2 (Infection Study) | 4.2 | 8 | 324 | 0.11 | 37% |
| Influenza, Fonville Study 3 (Vaccination Study, 1997) | 6.0 | 7 | 212 | 0.12 | 40% |
| Influenza, Fonville Study 4 (Vaccination Study, 1998) | 4.6 | 7 | 256 | 0.16 | 40% |
| Influenza, Fonville Study 5 (Vaccination Study, 2009) | 2.3 | 9 | 160 | 0.12 | 55% |
| Influenza, Fonville Study 6 (Vaccination Study, 2010) | 2.4 | 8 | 160 | 0.12 | 55% |
| HIV-1, Catnap (Monoclonal Antibodies) | 3.7 | 23 | 933 | (0.18) | (27%) |
| HIV-1, Catnap (Polyclonal Sera) | 3.1 | 7 | 71 | 0.19<br>(0.18) | 44%<br>(41%) |

**Figure S1.**
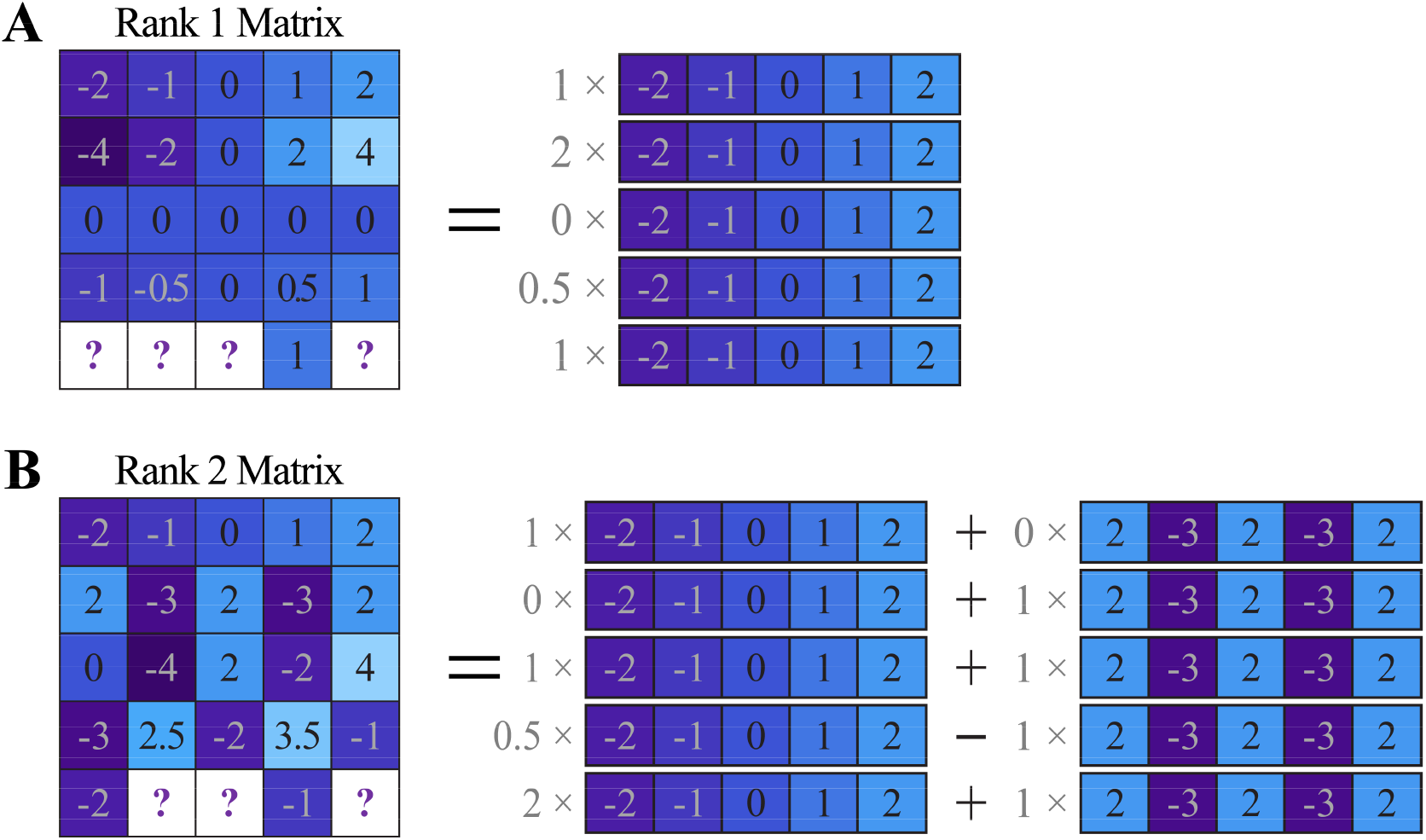
Example of low-rank matrices. (A) A rank 1 matrix has rows (or columns) that are multiples of a single vector. If the bottom row (representing a new antibody or virus) is added with just a single measurement, all other values in this row can be determined by finding the multiple of the common vector, as shown by the right-hand side of the equality. (B) A rank 2 matrix has rows (or columns) that are the linear combination of two vectors. If a new row is added with just two measurements, the other values in the row can be determined. Note that although the matrix on the left-hand side is visually very complex, it nevertheless follows a simple mathematical structure that can be determined using the singular value decomposition or alternative methods.

**Figure S2.**
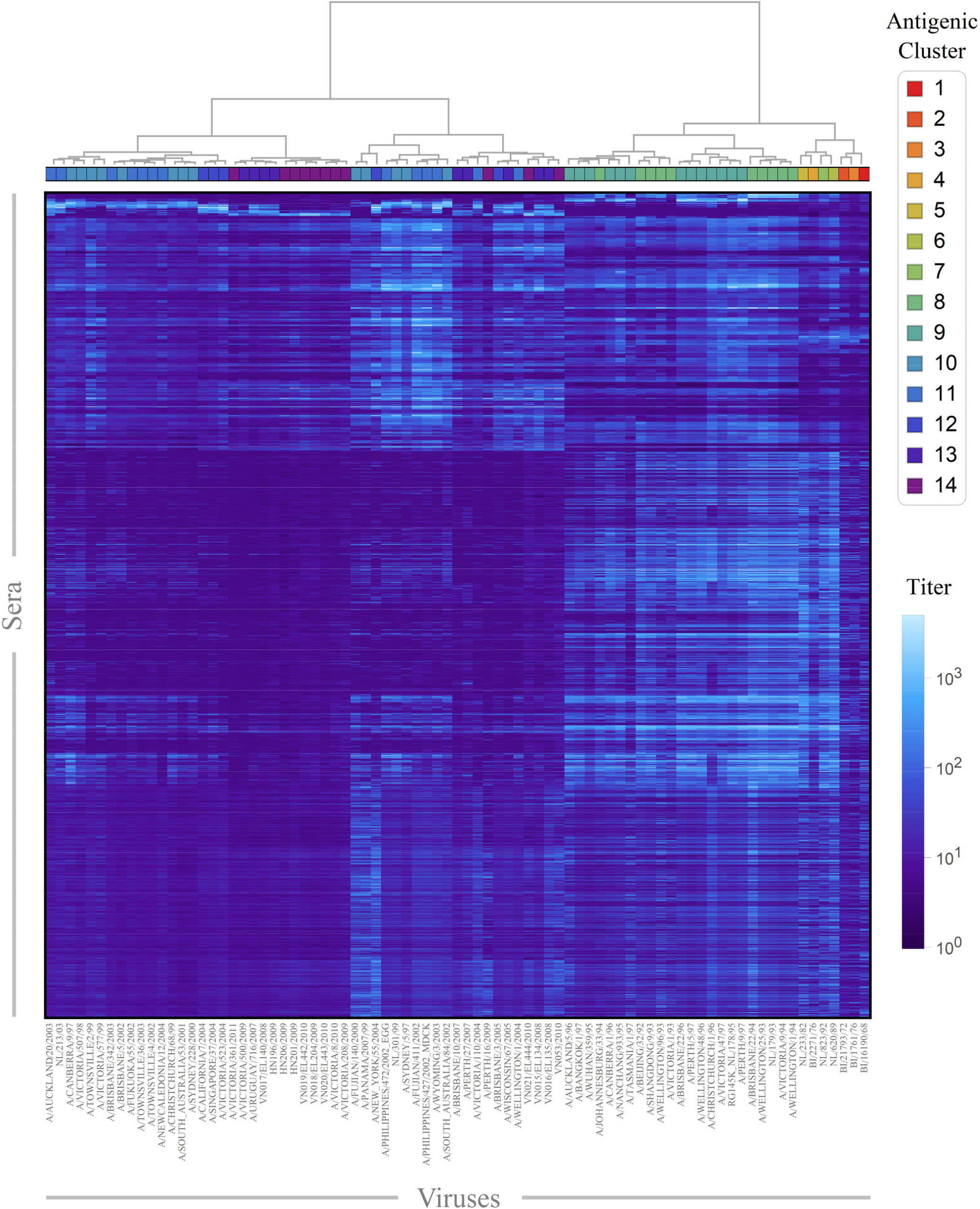
Antigenic clustering in the combined Fonville data. All six Fonville studies were combined, with any missing entries inferred by matrix completion. Heatmap values depict log_10_(IC_50_) for each virus-serum pair (columns and rows, respectively). Columns are labeled by antigenic cluster (defined in Figure S1 of Fonville 2014) and ordered according to hierarchical clustering of their profile across sera in the completed matrix using Ward’s minimum variance dissimilarity. Although many viruses group together by antigenic cluster, there are substantial differences within antigenic clusters, for example in antigenic clusters 10‒14.

**Figure S3.**
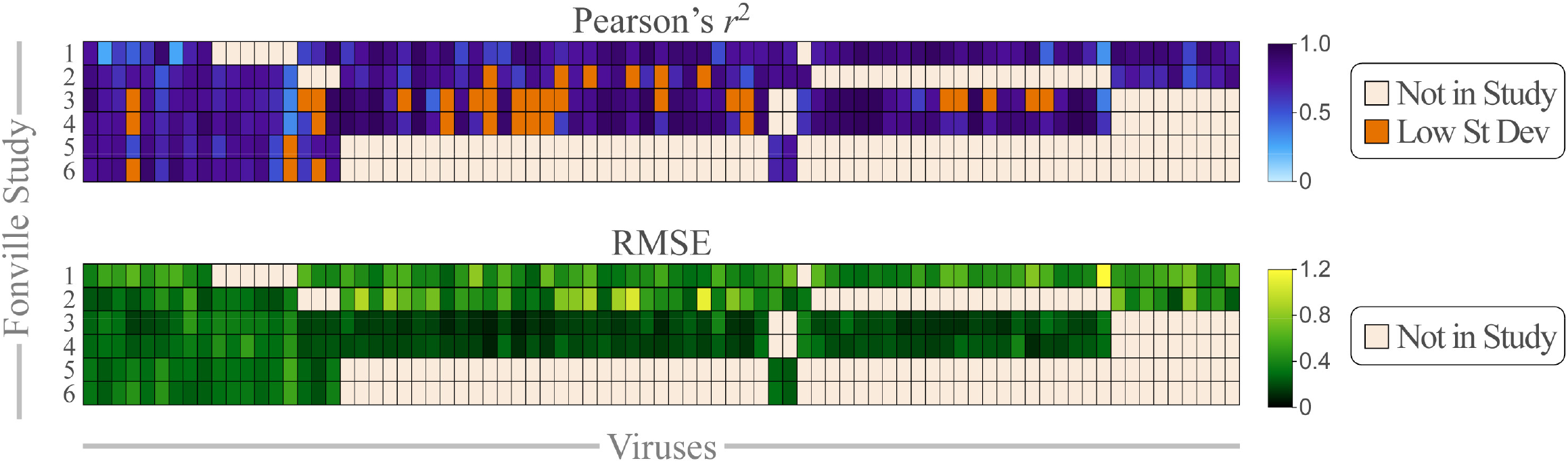
Inter-table completion using a single virus. As shown in Figure 4A,B, each virus was withheld from one Fonville study and predicted via matrix completion. Collectively, the measurements of the 81 viruses across the six Fonville studies led to 311 combinations. Pearson’s *r*^2^ [top] and the RMSE [bottom] are shown for each case. Viruses absent from a study are colored tan. Viruses shown in orange have predicted or measured log_10_(titers) with standard deviation ≤0.2; since they exhibit no linear dependence, their *r*^2^ is artificially deflated and they were excluded from the *r*^2^ analysis (see Methods).

**Figure S4.**
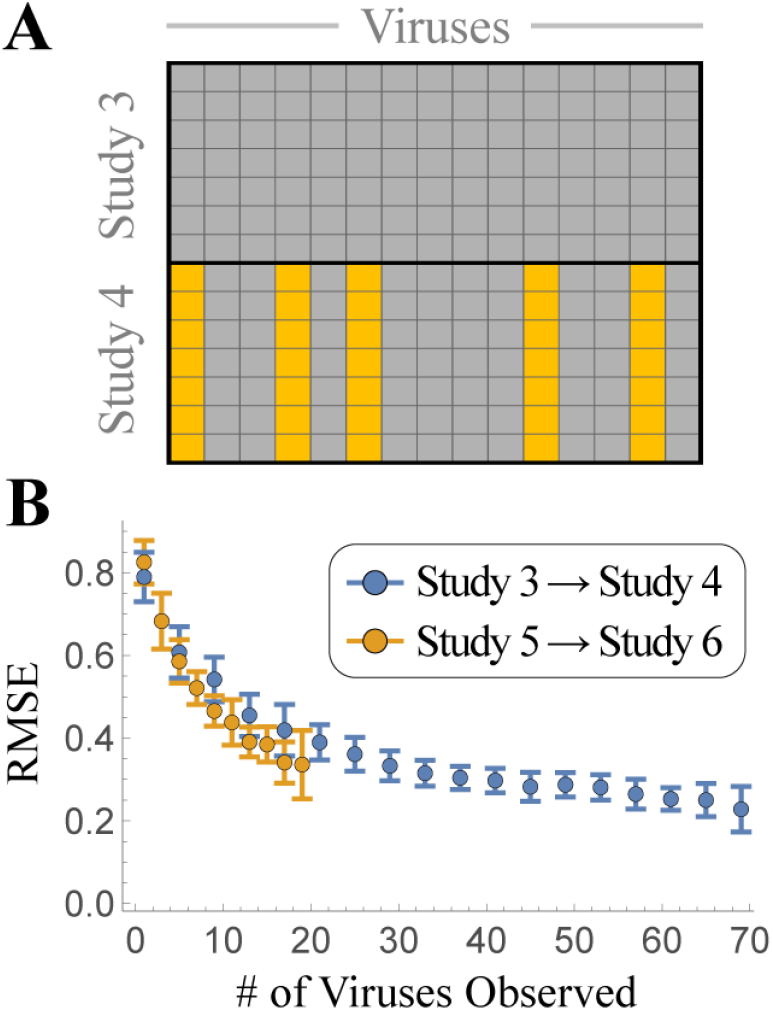
Inter-table completion using multiple viruses. (A) Fonville Studies 3 and 4 have identical virus panels with 70 viruses. Different subsets of viruses were entirely removed from Study 4 (*e.g*., five viruses shown in this schematic using columns of gold boxes) and inter-table completed. This process was repeated using Study 5 to inter-table complete Study 6, which contained identical virus panels with 20 viruses. (B) The number of viruses shown on the *x*-axis were observed in Study 4 or Study 6, and the remaining virus titers were imputed via matrix completion, and the resulting RMSE was calculated.

**Figure S5.**
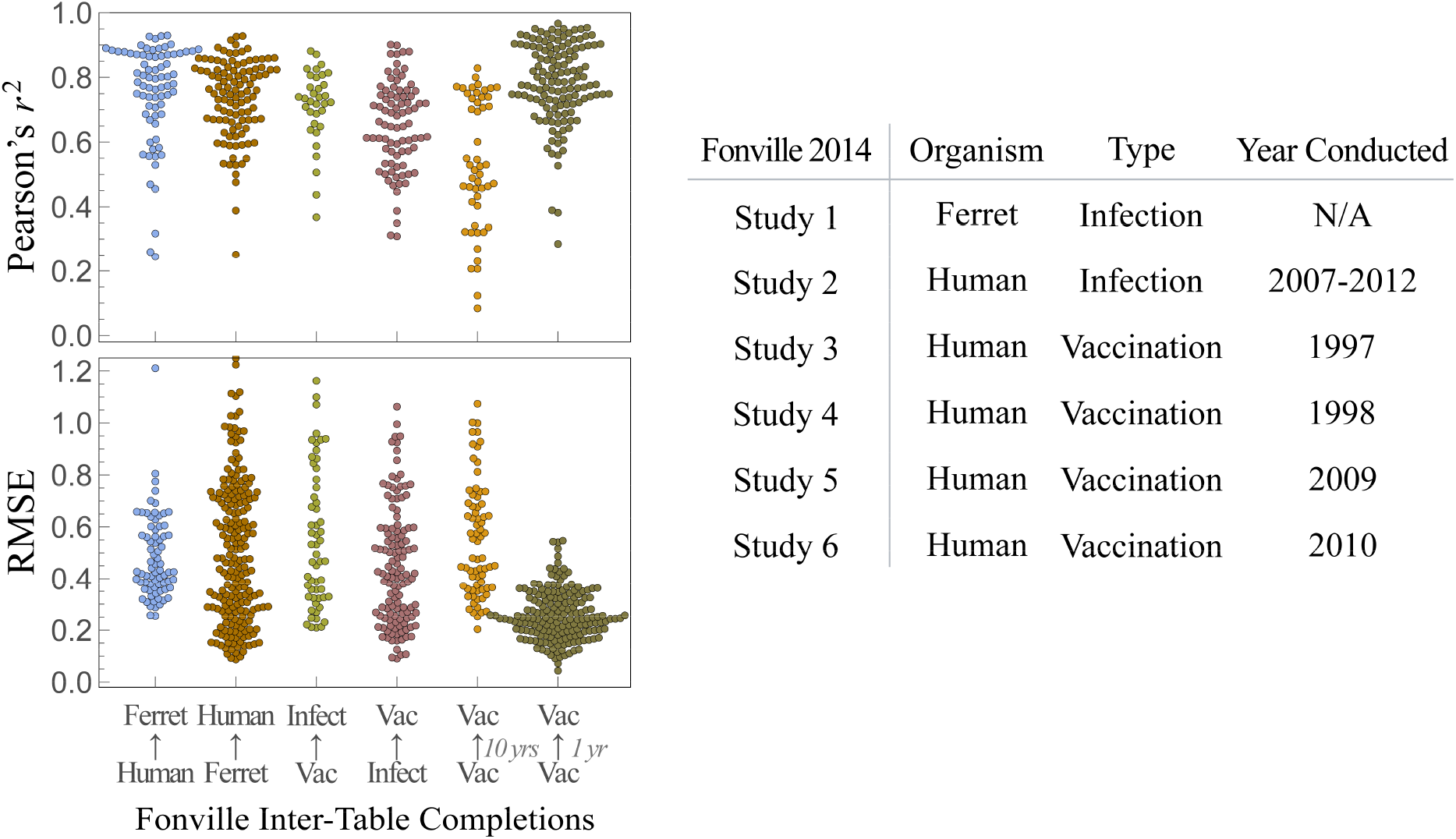
Inter-table completion across different types of data. Inter-table completion was performed analogous to Figure 4B, but using different subsets of Fonville studies. From left-to-right, the columns represents: (1) Using human data to impute ferret data [Studies 2-6 → Study 1] and (2) vice versa [Study 1 → Studies 2-6]; (3) using human vaccination data to impute human infection data [Studies 3-6 → Study 2] and (4) vice versa [Studies 2 → Study 3-6]; (5) inferring human vaccination studies using other human vaccination studies conducted 10 years apart [Studies 3-4 ↔ Studies 5-6]; and (6) inferring human vaccination studies using other human vaccination studies conducted 1 year apart [Study 3 ↔ Study 4 and Study 5 ↔ Study 6]. These results can be contrasted with the inter-table completion results in Figure 4B where the *j*^th^ column represents using *all* Fonville studies except Study *X* to impute the data in Study *X*. Hence, the first column in Figure 4B is identical to the left column in this figure [Studies 2-6 → Study 1]. The second column from the left in figure 4B [Studies 1,3,4,5,6 → Study 2] is similar to the third column in this figure, and comparing the two demonstrates the effects of excluding the ferret data from the inter-table completions. In general, including more datasets improves the inter-table completion results. The right-most column in this figure demonstrates that inter-table completion works best between vaccination studies that are 1 year apart, and this effect dominates the results in the four rightmost columns in Figure 4B.

**Figure S6.**
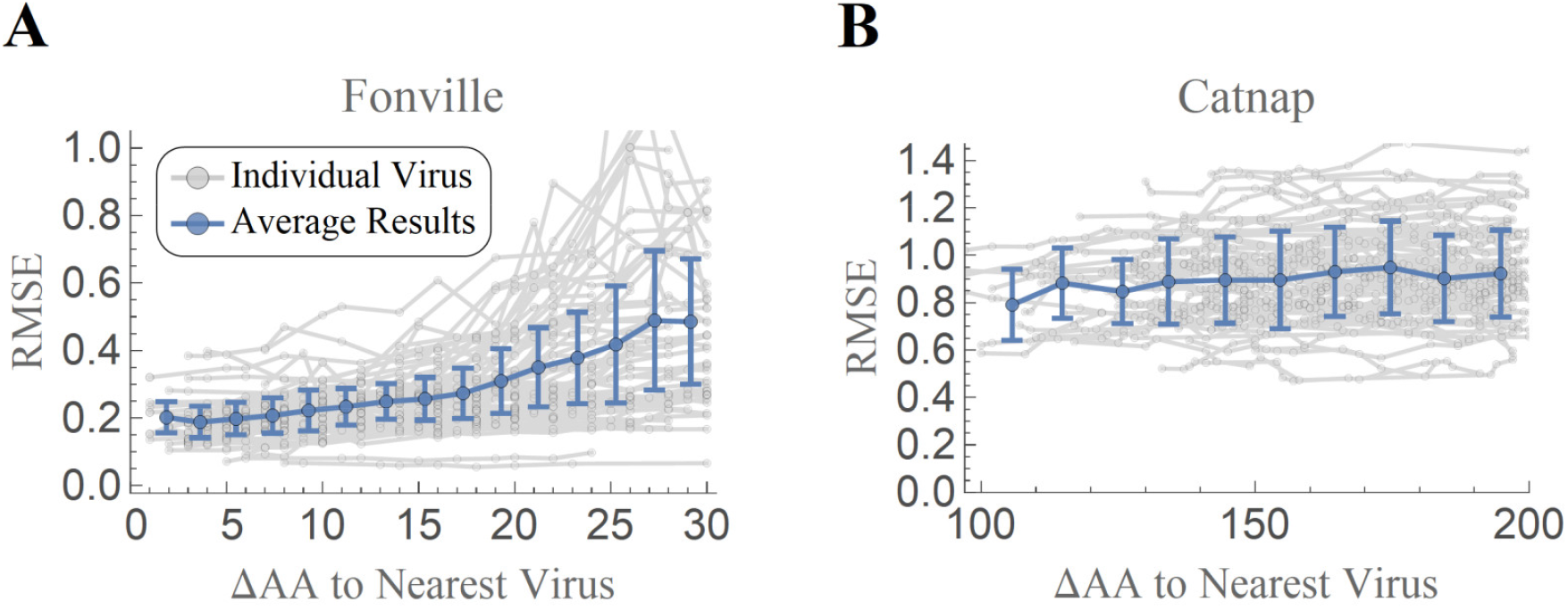
The accuracy of matrix completion diminishes for highly divergent viruses. The root-mean-squared error for viral titers as a function of the amino acid distance (ΔAA) to the most similar virus in a dataset. (A) For each virus *V* in Fonville Study 4 (chosen because it has nearly-complete data), construct a matrix containing the titers of *V* together with the titers of the 10 most similar viruses whose HA sequences are ≥ΔAA from *V*. Withhold 20% of titers from virus *V* and impute them via intra-table completion. Each gray line shows the resulting curve for one virus, and the blue line shows the average response over all viruses. The RMSE slowly increases from 0.16 when ΔAA≤5 (1.4-fold error) to 0.44 when ΔAA=30 (2.8-fold error). (B) This analysis was repeated for the more diverse Catnap viruses, except now using viruses whose Envelope sequence differed by ≥ΔAA. The response remained relatively flat with an RMSE≈0.9 between ΔAA=100 and ΔAA=200.

**Figure S7.**
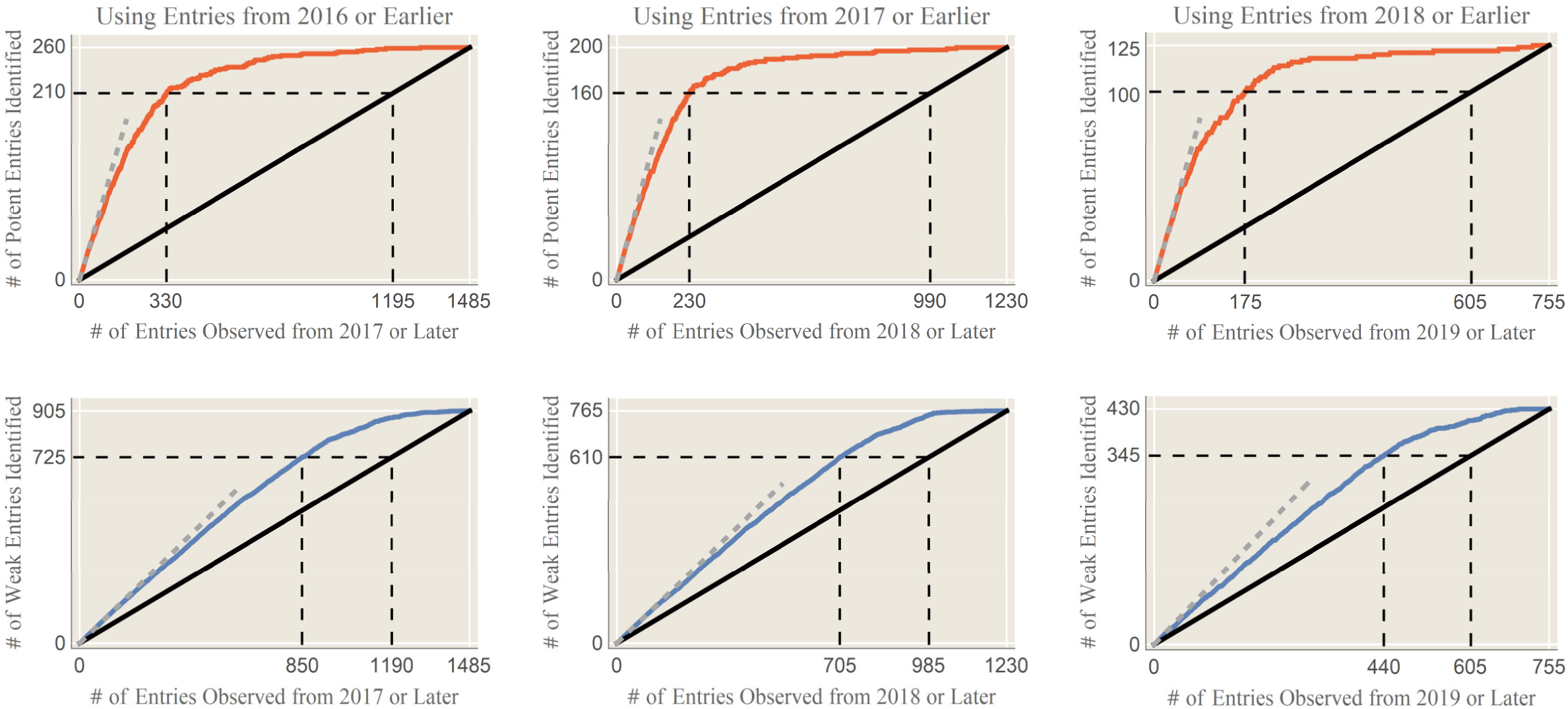
Using prior HIV-1 measurements in Catnap to predict future experiments. Matrix completion was applied to all antibody-virus measurements from 2016 [*left column*], 2017 [*middle column*], or 2018 [*right column*] to predict all future measurements. The predicted interactions were ordered from strongest-to-weakest [red curves in the *top row*] or weakest-to-strongest [blue curves in the *bottom row*], and each the true measurements were revealed in order. The top row shows the number of strong interactions measurements (IC_50_≤0.1 μg/mL) found after *N* measurements, either ordered by matrix completion or by a random search. The bottom row shows the analogous results when searching for weak interactions (IC_50_≥10 μg/mL). In all cases, matrix completion leads to a better discovery rate than a random search (black diagonal). The black dashed line shows when 80% of the strong/weak interactions are discovered. The gray dotted line (*y*=*x*) represents perfect predictions for the strong/weak interactions.

**Figure S8.**
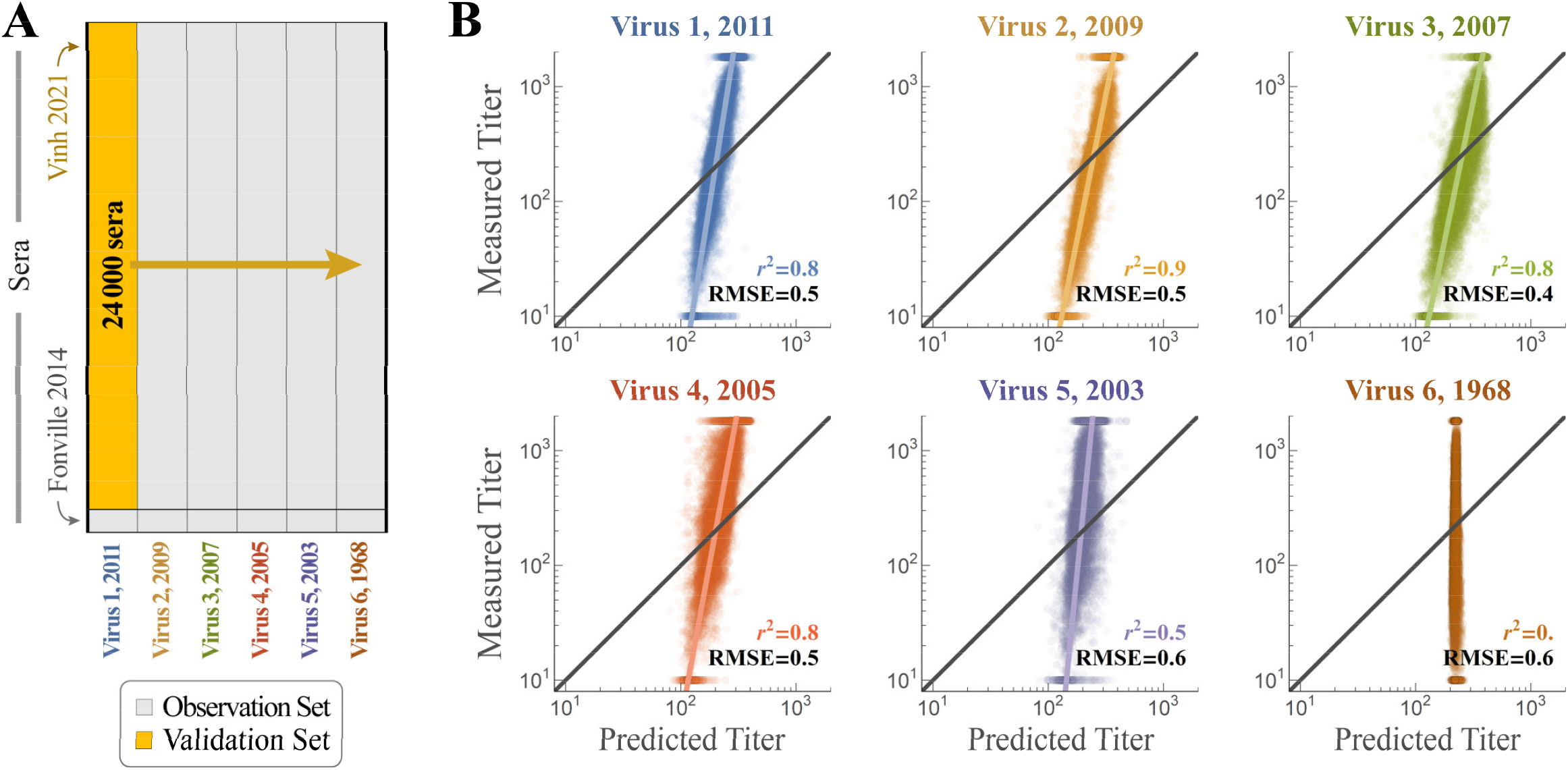
Using Fonville 2014 to Inter-Table Complete Vinh 2021. (A) 24,000 sera were measured against six H3N2 influenza viruses (Virus 1=A/Victoria/361/2011, Virus 2=A/Victoria/210/2009, Virus 3=A/Brisbane/10/2007, Virus 4=A/Wisconsin/67/2005, Virus 5=A/Wyoming/3/2003, Virus 6=A/Aichi/2/1968) (Nguyen Vinh et al., 2021). We remove all data from one of the Vinh strains and perform inter-table completion using completed measurements from the 1147 Fonville sera against these viruses. (B) The predicted versus measured titers when imputing the six different viruses. Although the predicted titers vary over a much smaller range of values (∼5-fold compared to the experimental measurements spanning 200-fold), Viruses 1-4 exhibit a strong correlation (*r*^2^≥0.8). Virus 6 serves as a negative control, demonstrating that a serum response to viruses from 2003-2011 cannot characterize the response against a virus from 1968.

## Notes

https://github.com/cleary-lab/Inhibition-Neutralization-Completion

### Competing Interest Statement

The authors have declared no competing interest.

